# The NAPstar family of NADP redox state sensors highlights glutathione as the primary mediator of anti-oxidative electron flux

**DOI:** 10.1101/2024.02.14.580349

**Authors:** Marie Scherschel, Jan-Ole Niemeier, Lianne J.H.C. Jacobs, Markus Hoffmann, Anika Diederich, Christopher Bell, Pascal Höhne, Sonja Raetz, Johanna B. Kroll, Janina Steinbeck, Sophie Lichtenauer, Jan Multhoff, Jannik Zimmermann, Tanmay Sadhanasatish, R. Alexander Rothemann, Carsten Grashoff, Joris Messens, Emmanuel Ampofo, Matthias Laschke, Jan Riemer, Leticia Prates Roma, Markus Schwarzländer, Bruce Morgan

**Author notes:** These authors contributed equally to this manuscript (Marie Scherschel, Jan-Ole Niemeier). To whom correspondence should be sent.

## Abstract

The NADPH/NADP^+^ redox couple is central to metabolism and redox signalling. NADP redox state is differentially regulated by distinct enzymatic machineries at the subcellular compartment level. Nonetheless, a detailed understanding of subcellular NADP redox dynamics is limited by the availability of appropriate tools. Here, we introduce NAPstars, a family of genetically encoded, fluorescent protein-based NADP redox state biosensors. NAPstars offer real-time, specific, pH-resistant measurements, across a broad-range of NADP redox states, with subcellular resolution. We establish NAPstar measurements in yeast, plants and mammalian cell models, revealing a conserved robustness of cytosolic NADP redox homeostasis. NAPstars uncovered NADP redox oscillations linked to the cell cycle in yeast and illumination- and hypoxia-dependent NADP redox changes in plant leaves. By selectively impairing the glutathione and thioredoxin anti-oxidative pathways under acute oxidative challenge, NAPstars demonstrated an unexpected role for the glutathione system as the primary mediator of anti-oxidative electron flux that is conserved across eukaryotic kingdoms.

## Introduction

Nicotinamide adenine dinucleotide phosphate (NADP), in its reduced (NADPH) and oxidized (NADP^+^) states, constitutes a central metabolic redox couple, which is found in all living organisms. NADPH plays a crucial role as an electron donor in numerous, typically anabolic, pathways, for example fatty acid and cholesterol synthesis and photosynthetic carbon assimilation, and for the enzymatic reduction of certain reactive oxygen species, including H_2_O_2_. NADPH also serves as the source of electrons for NADPH oxidases. This family of enzymes are central to the respiratory burst in immune responses and the generation of H_2_O_2_ as a second messenger in cellular signalling, development, and environmental sensing across kingdoms [1, 2].

Since its discovery in the early 1930s [3, 4], NADP, together with its cellular counterpart, NAD, has been extensively investigated. However, our understanding of subcellular NAD(P) redox dynamics is still remarkably incomplete, mainly due to a deficit of techniques allowing specific monitoring in defined subcellular compartments *in vivo*. Various crucial physiological parameters relate to pyridine nucleotide homeostasis including the concentration of the reduced, NAD(P)H, and oxidised forms, NAD(P)^+^. The final crucial parameter is the ratio between the two species, i.e. NADPH/NADP^+^ and NADH/NAD^+^, which enables the determination of the redox potential, i.e. *E*_NADP_ and *E*_NAD_. Indeed, *E*_NADP_ and *E*_NAD_ are parameters of central importance for cellular metabolism and regulation. Hereon, we use the terms NAD redox state and NADP redox state to refer to NADH/NAD^+^ (*E*_NAD_) and NADPH/NADP^+^ (*E*_NADP_) respectively.

Novel genetically encoded sensors allowed for accurate measurements of the subcellular dynamics of numerous metabolites, metals ions including calcium, H_2_O_2_, and pH [5–8]. Such sensors have revolutionised our understanding of countless previously intractable biological questions, particularly when dealing with small subcellular compartment-specific dynamics, and have underpinned numerous novel discoveries. Genetically encoded sensors for pyridine nucleotides were relative latecomers, however, significant progress has been made over the past decade in developing sensors for NADH [9], NAD^+^ [10, 11], NADH/NAD^+^ [12–14], NADPH [15], and NADP^+^ [16, 17]. However, existing NADPH sensors face major limitations, including sensitivity to pH, lack of responsiveness to NADP^+^, signal-to-noise ratio, and compatibility with alternative measurement techniques like FLIM. A recently created sensor, NERNST, was developed for monitoring NADP redox state [18]. Nonetheless, considerable concerns arise regarding the specificity of NERNST due to its dependence on a redox-sensitive green fluorescent protein (roGFP2) reporter, known to efficiently equilibrate with the glutathione redox couple *in vivo* [19–22].

In this study, we employed a rational probe design strategy to create the innovative NAPstar family of NADP redox state probes. The NAPstar family allows the monitoring of NADP redox states across a 5,000-fold range, spanning NADPH/NADP^+^ ratios from about 0.001 to 5. We showed that NAPstars facilitate specific and real-time monitoring of subcellular NADP redox state dynamics, which can be measured either by monitoring changes in fluorescence excitation and emission spectra or through fluorescence lifetime imaging. Applying NAPstars, we find a surprisingly oxidised cytosolic NADP redox state albeit with a remarkable robustness to oxidative challenges in yeast, human cells and plant. We used NAPstars to unveil oscillations of NADP redox state associated with cell division and metabolic cycles in yeast, as well as to monitor NADP redox dynamics influenced by illumination and hypoxia–reoxygenation in plants. Finally, we reveal a conserved, and predominant role of the glutathione system in mediating anti-oxidative electron flux in response to oxidative challenges across diverse eukaryotic cells.

## Results

We used the NAD redox state sensor, Peredox-mCherry (henceforth referred to as Peredox) [13], as a chassis for the development of novel NADP redox state probes (**Figure 1A**). Peredox incorporates a circularly permuted T-Sapphire (TS) fluorescent protein nestled between two copies of the NADH/NAD^+^-binding domain of the bacterial transcriptional repressor Rex [13]. Structural changes, contingent on whether the Rex domains are binding to NAD^+^ or NADH, induce changes in the TS fluorescence. This fluorescence change can be normalised against the signal from a C-terminally fused mCherry (mC) fluorescent protein. Peredox offers several advantages over other NADH and NADPH sensors, including pH-resistance and the apparent brightness of TS-based probes in biological systems relative to some cpYFP-based sensors like SoNar and the iNaps [12, 15].

**Figure 1.**
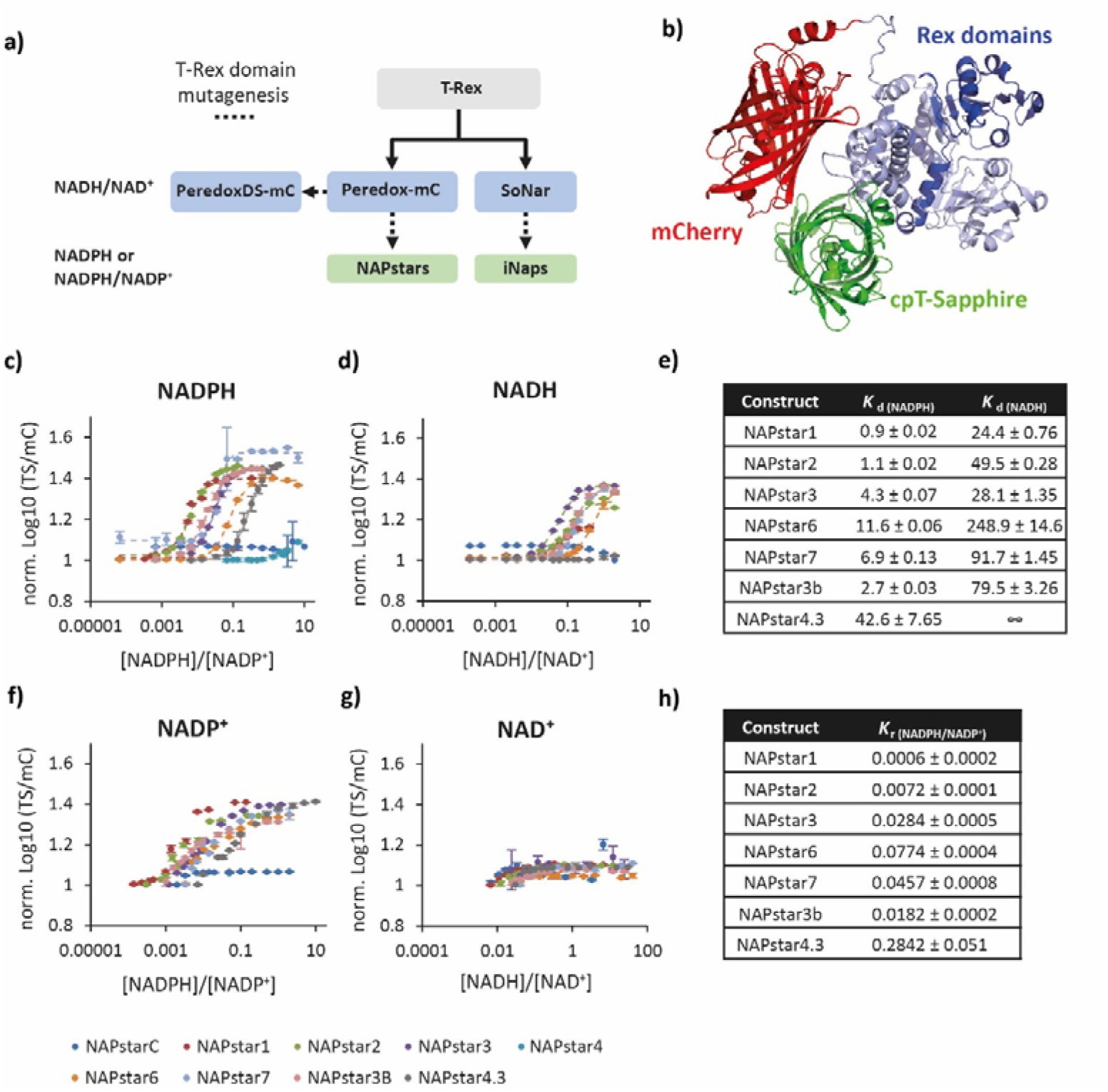
NAPstars respond specifically to changes in the NADP redox state. **a**, Diagram showing the development of selected NAD and NADP sensors including NAPstars. **b,** AlphaFold2 prediction of NAPstar structure. **c–d**, Graphs showing the normalised logarithm of the cpT-Sapphire/mCherry fluorescence ratio at **c**, different NADPH/NADP^+^ and **d**, NADH/NAD^+^ ratios. NADPH and NADH concentration was titrated against a fixed background of 150 µM NADP^+^ and 500 µM NAD^+^ respectively. In **c** and **d**, the dashed lines show a fitted sigmoidal function that was used to determine *K*_d(NAD(P)H)_. **e**, Table summarising the determined *K*_d(NADPH)_ and *K*_d(NADH)_ values of all NAPstars. **f–g**, Graphs showing the normalised logarithm of the cpT-Sapphire/mCherry fluorescence ratio at **f**, different NADPH/NADP^+^ and **g**, NADH/NAD^+^ ratios. NADP^+^ and NAD^+^ concentration was titrated against a fixed concentration of NADPH and NADH respectively that for each probe corresponded approximately to the determined *K*_d(NADPH)_ and *K*_d(NADH)_ values. **h**, Table summarising the determined *K*_r(NADPH/NADP+)_ for all NAPstars. For panels **c,d,f,g**, n=3 technical replicates. Data are presented as mean ± s.d.

To alter the binding pocket of Peredox to favour NADP binding, we introduced mutations known to switch the specificity of the Rex domain from favouring NADH binding to favouring NADPH [15] and generated combinatorial mutants thereof. These mutations were applied equally to each of the Rex domains of Peredox, resulting in a family of constructs termed NAPstars (**Table 1**). NAPstar1, 2, 3, 4, 6, 7 and NAPstarC were expressed as recombinant proteins in *E. coli* for *in vitro* characterisation. Fluorescence spectra were recorded in the presence of 0–100 µM NADPH (**Figure S1**). NAPstarC, containing mutations preventing nucleotide binding to Rex, remained unresponsive to changes in NADPH concentration (**Figure S1A,B**). NAPstars1, 2, 3, 6 and 7 exhibited pronounced NADPH-dependent changes in TS fluorescence excitation and emission spectra, with excitation and emission maxima at approximately 400 and 515 nm, respectively, and with a spectroscopic dynamic range of about 2.5, i.e. about 0.4 units on a log10 scale, similar to Peredox, (**Figure 1C**) [23]. Changes in NADPH concentration did not affect the fluorescence excitation and emission of mC (**Figure S1C–N**). Our results thus demonstrate that NAPstars are responsive to changes in NADPH concentration.

To gain insight into the potential structure of NAPstar, we employed AlphaFold2 with the 919 amino acids of Napstar3 as input. The sequence consists of a T-rex domain linked with an SAAGGH amino acid sequence to circular permutated TS, followed by a single amino acid Thr linked to a second T-rex domain, a GSGTGGNASDGGGSGG linker, and the mCherry sequence. Five models were generated, and the top-ranked model displayed a notably reliable structure, with an average Predicted Local Distance Difference Test (pLDDT) score of 87.8% (**Figure 1B**).

To better characterise the pyridine nucleotide specificity of the NAPstar constructs, we titrated NAPstar1, 2, 3, 6 and 7 with varying concentrations of NADPH, NADP^+^, NADH or NAD^+^ (**Figure 1C,D,F,G**). NADPH concentration was adjusted from 0.01–1000 µM in the presence of a constant 150 µM NADP^+^. For all constructs, except for the non-binding control NAPstarC and NAPstar4, we observed an NADPH concentration-dependent change in the TS/mC fluorescence emission ratio (**Figure 1C**). We determined apparent dissociation constants for NADPH (*K*_d(NADPH)_) from the fluorescence changes, ranging from 0.9 µM for NAPstar1 to 11.6 µM for NAPstar6 (**Figure 1E**). All NAPstars showed affinity to NADH, titrated in the presence of 500 µM NAD^+^, with *K*_d(NADH)_ ranging from 24.4–248.9 µM. However, affinity to NADH is substantially weakened compared to Peredox (*K*_d(NADH)_ 1.2) [23] and for each NAPstar is one to two orders of magnitude lower than for NADPH (**Figure 1D,E**). For all NAPstars, we also observed ratiometric fluorescence responses to NADP^+^ in the opposite direction to those induced by NADPH (**Figure 1F**). Such changes were not observed for NAD^+^ (**Figure 1G**). These results thus suggest that NAPstars report the *bona fide* NADP redox state rather than responding solely to NADPH concentration. To further investigate this, we monitored the dependence of the TS/mC ratio on the NADPH/NADP^+^ ratio at different total NADP (NADPH + NADP^+^) pool sizes of 100, 300 and 500 µM respectively (**Figure S2**). The responses of NAPstar sensors remained largely stable across various pool sizes, indicating a predominant sensitivity to the NADP redox state rather than the individual concentrations of NADPH or NADP^+^. Exceptions seem to be NAPstar6 and 7, which displayed some dependence on pool size. Those two variants also exhibited the highest *K*_d(NADPH)_. To reflect the measurement of the NADPH/NADP^+^ ratio, rather than the NADPH concentration, from now on we report *K*_ratios_ (*K*_r(NADPH/NADP+)_) for every NAPstar (**Figure 1H**).

### Mixing different Rex domains to enhance control of NAD(P)H binding affinity

NAPstars contain both the Rex domains for NADP binding. In contrast, iNaps presumably require dimerisation to form the Rex-dimer for a functional sensor unit, adding sensor concentration (i.e. expression levels) as an additional variable [15]. We asked if we could exploit the fact that NAPstars encompass a Rex dimer within one polypeptide, by introducing a blend of two different Rex domains into one NAPstar sensor, with the aim to expand the range of accessible NADP redox states. As a first attempt, we created NAPstar3b, featuring a single point mutation, V126Y, in the N-terminal NAPstar3-derived Rex domain instead of the usual V130Y mutation which was kept in the second, C-terminal, NAPstar3-derived Rex domain. NAPstar3b was found to have a slightly decreased *K*_d(NADPH)_ but an approximately 3-fold higher *K*_d(NADH)_ (**Figure 1C–H**). Building on this idea, we further pushed the boundaries by combining an N-terminal Rex domain from NAPstar4, a NAPstar variant initially dismissed due to lack of detectable affinity for NADPH (**Figure 1C**), with a C-terminal Rex domain from NAPstar3. The resulting construct, NAPstar4.3, was purified as a recombinant protein for *in vitro* characterisation (**Figure 1C–H**). Intriguingly, we found that NAPstar4.3 had a strongly decreased affinity for NADPH, *K*_d (NADPH)_ = 42.6 µM, *K*_r(NADPH/NADP+)_ = 0.28 (**Figure 1C,E**), and no detectable affinity for NADH (**Figure 1D,E).** This suggests that combining differently mutated Rex domains within a single NAPstar construct provides enhanced control to rationally modulate NAPstar binding properties.

### NAPstars are pH resistant

Resistance of the fluorescence excitation and emission spectra to pH changes is a major achievement of the molecular engineering of the Peredox probe (**Figure S3A–H**) [13] and there is no reason to believe that this characteristic would differ in the case of NAPstars. To verify whether the NAPstars do indeed retain the pH-resistance seen for Peredox, we incubated each NAPstar probe in various buffers with pH ranging from 6.0–9.0. Each NAPstar was incubated in the absence of NADPH and NADP^+^, in the presence of a high NADPH/NADP^+^ ratio mixture, and in the presence of a low NADPH/NADP^+^ ratio mixture. The TS/mC ratio of every NAPstar varied by only ∼30% over a 3 pH-unit range. The change in TS/mC when NAPstars were incubated with high NADPH/NADP^+^ ratio mixtures was even lower, remaining below 10% for all NAPstars. In summary, NAPstars show remarkable resistance of their fluorescence response to changes in pH.

### NAPstars specifically respond to dynamic NADP redox changes in vitro

To further test the dynamic and specific responsiveness of NAPstars to changes in NADP redox status, we used recombinant NAPstar probes to monitor NADP dynamics in enzyme assays *in vitro* (**Figure S4**). First, we monitored the reduction of NADP^+^ to NADPH in the isocitrate dehydrogenase-catalysed reaction from isocitrate to α-ketoglutarate (**Figure S4A)**. Upon the addition of isocitrate to start the reaction, NAPstar1, 2, and 3 exhibited a rapid increase in the TS/mC ratio. The response rate correlated with the *K*_r(NADPH/NADP+)_, i.e. NAPstar1 and 2 responded most rapidly, followed by NAPstar3. No response was detected for NAPstarC. As a further control for the conversion of NADP^+^ to NADPH, we simultaneously detected an increase in NADPH autofluorescence after initiating the reaction.

Next, we monitored NADP oxidation, i.e. NADPH to NADP^+^ conversion during the glutathione reductase-catalysed reaction between NADPH and GSSG (**Figure S4B**). Upon the addition of GSSG, we observed a rapid decrease in the TS/mC ratio for each NAPstar construct except NAPstarC. NAPstar3, with the highest *K*_r(NADPH/NADP+)_ among the tested NAPstars, responded first to NADPH consumption, followed by NAPstar1 and 2.

Finally, we used the NAPstars and Peredox, to monitor the conversion of NADPH to NADH, which cannot be resolved using standard NAD(P)H autofluorescence (**Figure S4C**). Upon initiating the reaction with alkaline phosphatase, we observed a decrease in the TS/mC ratio, first for NAPstar3, followed by NAPstar2, and finally NAPstar1. The TS/mC of Peredox increased rapidly following the initiation of the reaction, consistent with an increase in NADH concentration. In summary, these results demonstrate that NAPstars and Peredox allow selective monitoring of NADP and NAD dynamics, respectively.

### NAPstars are excellently suited to FLIM measurements

Fluorescence lifetime imaging (FLIM) is a powerful alternative approach to monitor the status of fluorescent molecules and proteins and can be especially advantageous, for example in situations of low signal-to-noise or where there are overlapping fluorescence excitation and emission spectra. We thus sought to test the suitability of NAPstars sensors to FLIM measurements (**Figure S5)**. To this end, we monitored the fluorescence lifetime of NAPstar4.3 in the presence of different NADPH/NADP^+^ ratios. We observed a change in fluorescence lifetime from 1.3 ns in an NADP^+^-saturated state to 2.3 ns in an NADPH-saturated state, which is exceptionally large for a fluorescent protein-based biosensor. Based on the FLIM data, we determined a *K*_d(NADH)_ = 40.5 µM and *K*_r(NADPH/NADP+)_ = 0.27, which are very close to the 42.6 µM and 0.28 values determined by the fluorescence intensity measurements (**Figure 1**). We conclude that NAPstars have favourable characteristics for the development of FLIM-based measurement approaches.

### Cytosolic NADP redox homeostasis is robustly maintained in yeast

Confident that NAPstars are indeed specific sensors of the NADP redox state *in vitro*, we next sought to test them in various eukaryotic model systems, starting with the budding yeast, *Saccharomyces cerevisiae*. We used a microplate-based assay to monitor the response of all NAPstars and PeredoxDS (an affinity variant of Peredox-mCherry for more reducing NAD redox states) [23], expressed in the yeast cytosol, to exogenous H_2_O_2_ at an initial concentration ranging from 0–5 mM (**Figure 2** and **Figure S6**). Strikingly, even at the highest H_2_O_2_ concentration, minimal response was detected for NAPstar1, 2 or 3 (**Figure 2C** and **Figure S6**). NAPstar6 and 7, with a high *K*_r(NADPH/NADP+)_, showed a limited response with a rapid recovery, while the TS/mC ratio of NAPstar4.3 was close to that of NAPstarC (**Figure S6**). These observations are consistent with the relative affinities of the sensor variants determined *in vitro* (**Figure 1**) and suggest robust and active maintenance of NADP redox state even under severe oxidative challenge, which is in stark contrast to previous observations of a highly volatile cytosolic NAD redox state as monitored by Peredox [13, 23]. The similarity in the fluorescence ratio of NADP^+^-bound and unbound NAPstars (as for NAPstarC) is consistent with previously published crystal structures of *Thermotoga maritima* Rex in NADH-bound, NAD^+^-bound, and unbound (apo) states [24]. Unbound and NAD^+^-bound TmRex are structurally very similar, while NADH binding induces significant structural changes. This suggests that NAPstar4.3 is in an almost fully NADP^+^-bound state in the yeast cytosol, unable of responding to a further oxidation of the NADP redox couple (**Figure S6**). The measurements further allow a first *in vivo* estimate for the NADPH/NADP^+^ ratio in the cultured yeast cytosol in the range of 0.03–0.3 (**Figure 1, 2 and Figure S6**).

**Figure 2.**
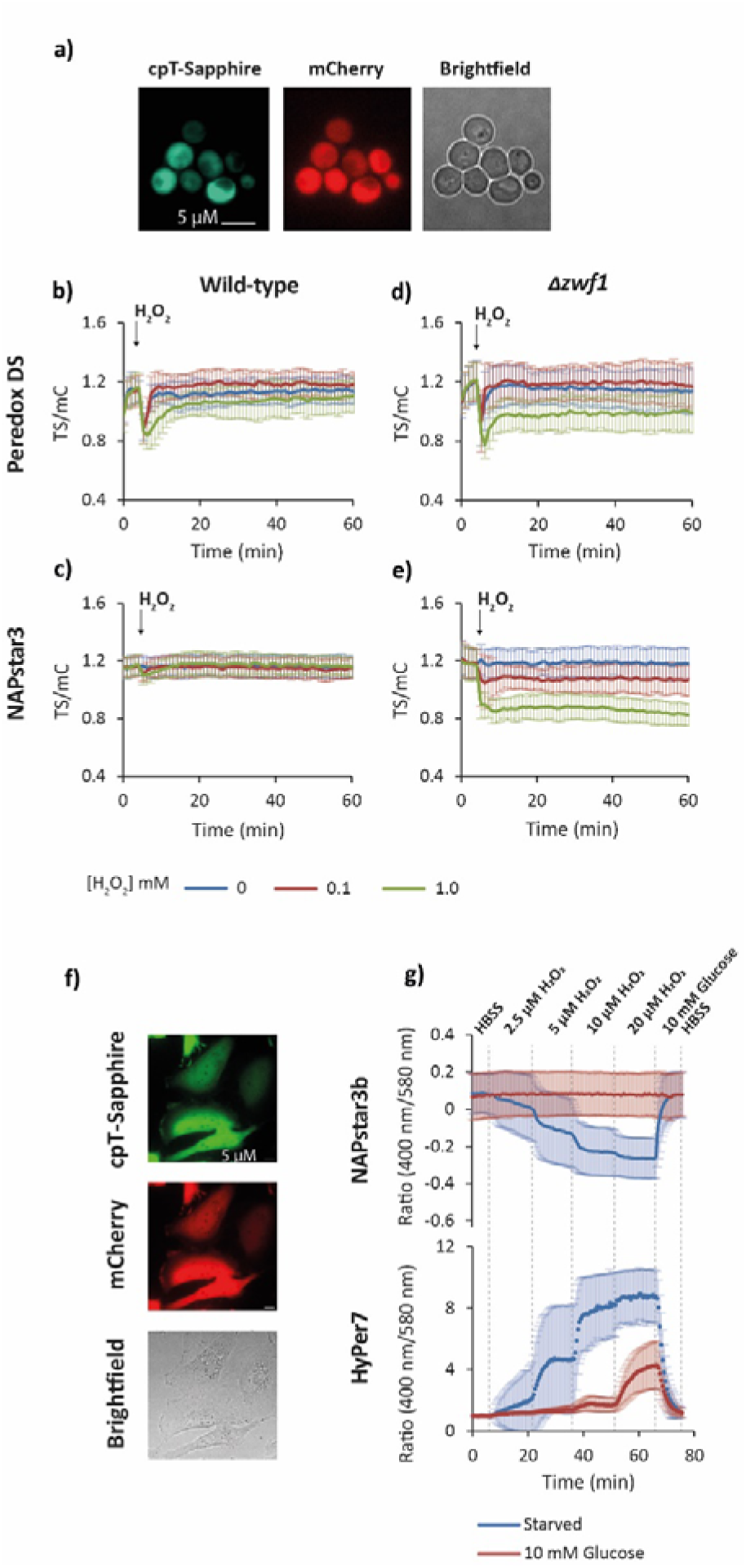
Cytosolic NADP redox homeostasis is robustly maintained. **a**, Epifluorescence microscopy images of yeast cells expressing NAPstar3, showing cpT-Sapphire and mCherry fluorescence as well as brightfield microscopy. **b–e,** Response of PeredoxDS and NAPstar3 probes, expressed in the cytosol of wild-type and Δ*zwf1* yeast cells, to the addition of exogenous H_2_O_2_ at the indicated concentrations (n=3 experimental repeats in which H_2_O_2_ responses was monitored in cells derived from independent cultures). **f**, Epifluorescence microscopy images showing cpT-Sapphire and mCherry fluorescence of NAPstar3b expressed in the cytosol of HeLa cells. **g,** Response to NAPstar3b (n=151 individual cells in total, monitored in the course of four experimental replicates) and HyPer7 (n=154 individual cells in total, monitored in the course of three experimental replicates) probes, expressed in the cytosol of HeLa cells cultured in a perfusion chamber, to the perfusion of buffers with stepwise increases in H_2_O_2_ concentration. Data are presented as mean ± s.d.

To distinguish between *bona fide* robustness of cytosolic NADP redox maintenance and a potential lack of probe functionality in yeast, we constrained cytosolic NADPH regeneration. To this end, we monitored the NAPstar responses in a Δ*zwf1* yeast strain. *ZWF1* encodes glucose 6-phosphate dehydrogenase, the first enzyme in the pentose phosphate pathway, which is the major source of cytosolic NADPH in yeast. In Δ*zwf1* cells, we observed sensitive, H_2_O_2_ concentration-dependent responses for all NAPstar constructs, supporting the conclusion that cytosolic NADP redox homeostasis is robustly maintained in wild-type yeast by adjusting flux through the oxidative pentose phosphate pathway (**Figure 2** and **Figure S6**). This observation is reminiscent of our previous observations of the robustness of cytosolic glutathione redox homeostasis, which largely relies on robust NADPH supply, indicating that the robustness in NADP redox state also serves the stability of downstream redox pools [21].

### Robustness of cytosolic NADP redox homeostasis is conserved

We next sought to understand to what extent the principles of NADP redox maintenance apply beyond yeast. Hence, we introduced and validated NAPstars in additional eukaryotic models. To this end, we used transient transfection to introduce NAPstar3b into HeLa cells (**Figure 2F**). We were first interested in monitoring the response of NAPstar3b to exogenous H_2_O_2_ in cells subjected to glucose starvation, comparing them to cells continuously exposed to 10 mM glucose. In a perfusion chamber, we monitored the response of HeLa cells to stepwise increases in H_2_O_2_ concentration (**Figure 2G**). In glucose-starved cells, we observed a stepwise increase in NADP oxidation. In contrast, in non-glucose starved cells, with the presence of 10 mM glucose in all perfusion buffers, no NADP oxidation was observed, even with continuous perfusion of buffer containing 20 µM H_2_O_2_. Additionally, we used the ultra-sensitive H_2_O_2_ probe, HyPer7 [25], to monitor H_2_O_2_ in the same experimental setup (**Figure 2G**). Consistent with the NAPstar response, we observed almost no HyPer7 response in glucose-fed cells, except at the highest H_2_O_2_ concentration, suggesting that HeLa cells can very efficiently remove H_2_O_2_. In contrast, in glucose-starved cells, we observed a strong HyPer7 response beginning with the lowest H_2_O_2_ concentration tested.

In summary, we find that, akin to yeast, cytosolic NADP redox homeostasis in HeLa cells is robustly maintained, and NADPH is readily available for the reduction of exogenous H_2_O_2_. Robustly regulated cytosolic NADP redox homeostasis hence may be a conserved cell physiological characteristic from yeast to human cells.

### Yeast cell cycle is accompanied by oscillations in NAD and NADP redox state

Confident that NAPstars and Peredox are faithfully reporting changes in NADP and NAD redox states within the yeast cytosol, we tested whether we could use NAPstars and Peredox to investigate the existence of dynamic changes in pyridine nucleotide redox state during the yeast metabolic cycle (YMC). The YMC is a phenomenon observed in continuous yeast cultures under mild glucose limitation, involving synchronised metabolic and transcriptional cycles along with cell cycle synchronisation [26, 27]. Previous reports have suggested that the total NAD(H) and NADP(H) levels change during the YMC [28], or have shown changes in NAD(P)H [29]. However, measurement of changes in NADP, or NAD redox states, and examining specific subcellular NAD(P) pools has hitherto been impossible in this system.

To address this, we established continuous yeast cultures [30] with yeast cells expressing Peredox, NAPstar4.3, or HyPer7 [25]. After testing different NAPstars we chose NAPstar4.3 for the specific conditions of the fermentor cultures, where this sensor variant showed optimal performance, unlike in more nutrient-rich batch cultures (**Figure 3 and Figure S7**). Using a coupled fermentor–fluorimeter setup to continuously monitor probe fluorescence within the cells in the fermentor culture (**Figure 3A,B**) [30], we found periodic oscillations in both NAD and NADP redox states (**Figure 3C**). These oscillations were synchronized with well-established fluctuations in oxygen consumption and to cell division. HyPer7 responses confirmed our previously reported cell cycle and metabolic cycle-associated H_2_O_2_ cycles, based on roGFP2-Tsa2ΔC_R_ sensor responses [30, 31]. Interestingly, our measurements showed that the three different redox species responded independently of each other, are not in equilibrium, and need to be analysed separately. For example, the peak reduction of the NADP pool coincided with the highest H_2_O_2_ levels. Likewise, the reduction and oxidation of the NAD and NADP pools occurs independently. These experiments highlight the complex and highly dynamic redox landscape of the cell, where individual redox couples, even within a single subcellular compartment, often do not equilibrate with each other *in vivo*. Those data re-emphasize the fact that there is no such thing as a general cellular redox state and the critical need for the specific empirical measurement of individual redox couples.

**Figure 3.**
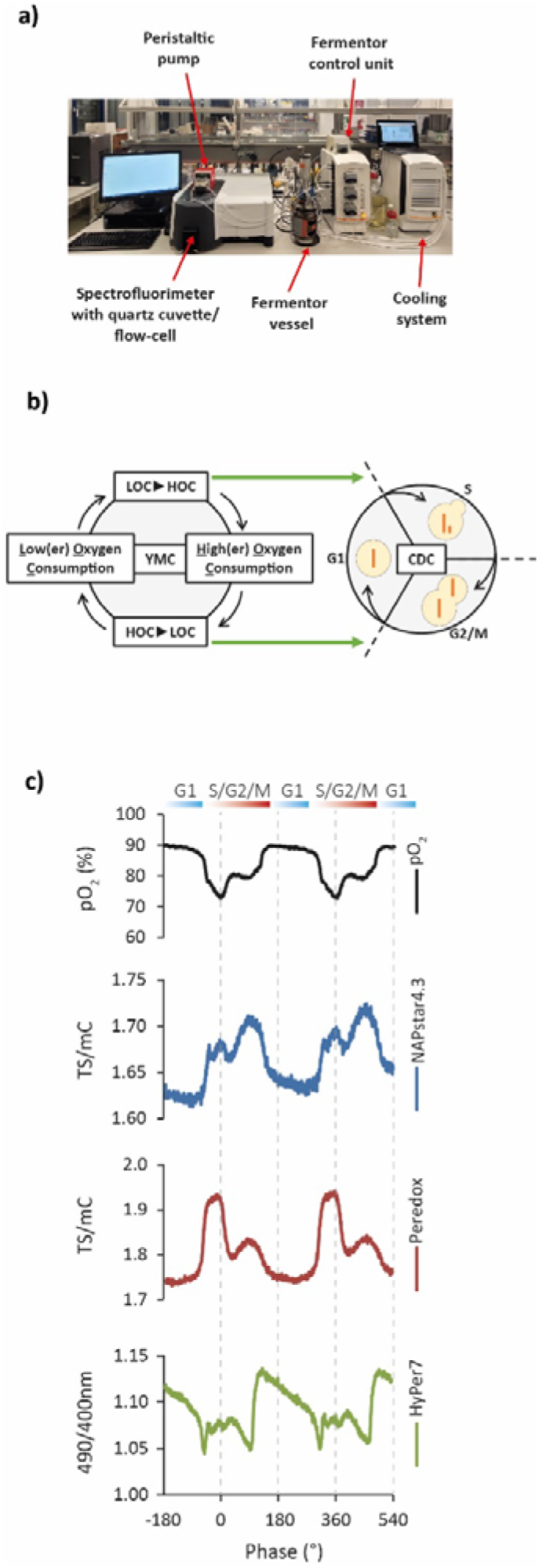
Oscillations in cytosolic NADP and NAD redox state accompany the yeast metabolic cycle (YMC). **a,** Photograph of the coupled fermentor–fluorimeter setup used in our lab to monitor redox changes in YMC-synchronised cultures. **b,** Diagram illustrating the coupled metabolic and cell division cycles observed during the YMC. CDC (cell division cycle), HOC (high oxygen consumption) and LOC (low oxygen consumption. **c**, Representative traces showing the changes in dissolved oxygen, NAPstar4.3 (NADP redox state), Peredox (NAD redox state) and Hyper7 (H_2_O_2_) during two complete cycles of the YMC (n=2, in which probe dynamics were measured for multiple YMC cycles in two independent YMC-synchronised cultures; **Figure S7**).

### NAPstars reveal NADP redox responses in the plant cytosol to illumination and hypoxia

Next, we aimed to enable NAPstar-based measurements of the NADP redox state in plants, which has a crucial role in underpinning photosynthesis and stress responses. Despite its importance in crop improvement, the specific dynamics of compartmentalized NADP redox state have been difficult to assess [32–34]. Using *Arabidopsis thaliana* as a model, we generated stable transgenic NAPstar3, NAPstar4.3 and NAPstarC lines with cyto-nuclear sensor expression. (**Figure 4A** and **Figure S8**). The steady-state TS/mC ratio of NAPstar3 was higher than for NAPstar4.3 reflecting the lower *K*_r(NADPH/NADP+)_ of NAPstar3. Consistent with our observations in yeast, the difference in NADPH occupancy suggests a close match of the response ranges of both sensor variants with the physiological NADP redox state (**Figure S8**), which may be estimated at about 0.3.

**Figure 4.**
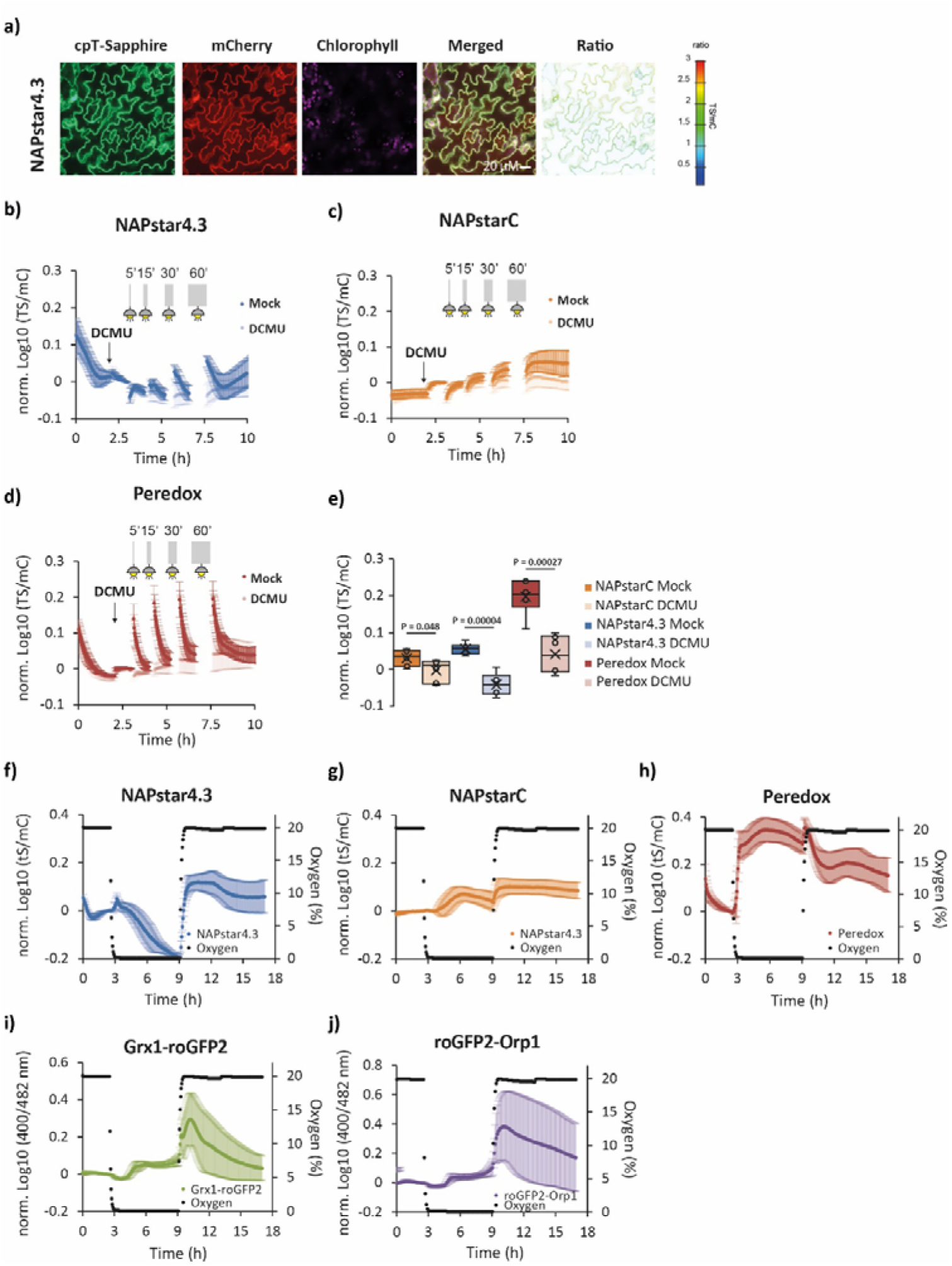
Cytosolic redox dynamics accompany hypoxia-reoxygenation and illumination in plants. **a,** Confocal microscopy images of NAPstar4.3 expressed in the cytosol of *Arabidopsis thaliana* plants. Scale bar = 20 µM. **b–d,** Response of NAPstar4.3 (**b**), NAPstarC (**c**) and Peredox (**d**) to the indicated periods of illumination after treatment with the photosynthetic inhibitor DCMU or a vehicle control (n=6 leaf discs from individual plants). **e,** Box and whisker plot showing the change in the normalised log10 TS/mC ratio after 60 minutes of illumination, P-values are derived from an unpaired two-tailed Student’s *t*-test. **f–j,** Response of NAPstar4.3 (**f**), NAPstarC (**g**), Peredox (**h**), Grx1-roGFP2 (**i**) and roGFP2-Orp1 (**j**) probes to 6 hours of hypoxia (0.1% oxygen) followed by restoration of normal atmospheric oxygen levels (n=6–7 leaf discs from individual plants). In all panels, data are presented as mean ± s.d.

To assess the *in vivo* responsiveness of the sensors, we illuminated leaf tissue, reasoning that activation of photosynthesis triggers the exchange of redox equivalents between the chloroplast stroma and the cytosol by different metabolite shuttles (**Figure 4**) [35]. Consecutive illumination of discs from mature Arabidopsis leaves, for periods of 5, 15, 30 and 60 min, each of which was followed by fluorimetric time-lapse measurements, revealed a pronounced reduction of the cytosolic NAD pool as monitored by Peredox (**Figure 4D**) [23]. The amplitude of NAD reduction increased slightly with the duration of illumination (**Figure 4D**). In contrast, NAPstar4.3 did not show cytosolic NADP reduction after 5 and 15 min of illumination (**Figure 4B**). A reduction was observable only after the longer illumination periods, i.e. 30 and 60 min (**Figure 4B**). The amplitudes of the NAD(P) redox dynamics were diminished by the photosynthetic inhibitor 3-(3,4-dichlorophenyl)-1,1-dimethylurea (DCMU) (**Figure 4B–E**), validating that the dynamic responses were driven by photosynthetic electron transport. The NAPstarC control showed changes too, indicating an NADP-independent effect in leaf tissue. Those changes were however minor, compared to that of NAPstar4.3, and may be corrected for by subtraction (not performed here in the interest of full data transparency) (**Figure 4B,C**). Based on the observation that NAPstar4.3 allowed dynamic measurements of both reductive and oxidative changes in the cytosolic NADP pool, we selected NAPstar4.3 as the most suitable sensor variant for further measurements in plant leaf tissue.

We next set out to explore NADP redox dynamics in response to hypoxia stress, which causes a cellular redox crisis. Hypoxia-dependent, inhibition of respiration leads to NAD reduction, as previously monitored in living leaf tissue [36]. The impact of hypoxia on cytosolic NADP redox dynamics had remained less clear, since lack of oxygen as an electron sink causes metabolic reduction, but also boosts H_2_O_2_ production and glutathione oxidation in different phases of the hypoxic episode [37]. Lowering oxygen levels to 5%, 1% and 0.1% for 6 h before re-oxygenation to ambient levels confirmed fast and reversible NAD reduction in Arabidopsis leaf tissue at 1 and 0.1% oxygen (**Figure 4F–J and Figure S9**). Interestingly, NADP redox state, as detected by NAPstar 4.3, responded oppositely to NAD redox state. After a lag phase, the NADP pool was gradually oxidized. At the technical level this response validates the strict specificity of NAPstar4.3 for the NADP, and not NAD, redox state *in planta*. At the physiological level, it shows that the redox dynamics of the NADP and NAD pools are strictly independent in the plant cytosol, which is in line with the absence of any transhydrogenase homologue in the plant genome.

We next hypothesized that NADP oxidation during hypoxia may be due to NADPH consumption by the anti-oxidant machinery which is not matched by the rate of re-supply due to limitations in flux through central metabolism. To test this hypothesis, we also measured glutathione redox potential and H_2_O_2_ dynamics using Grx1-roGFP2 and roGFP2-Orp1, respectively. Both sensors showed increased oxidation during 1% and 0.1% hypoxia, which correlated with NADP oxidation, suggestive of efficient equilibration between NADP and glutathione redox status via glutathione reductase in the plant cytosol. This correlation was lost upon re-oxygenation, however, as the NADP pool was rapidly reduced, while both Grx1-roGFP2 and roGFP2-Orp1 reported a burst in oxidation (**Figure 4F–J**). This observation might be explained by a burst of H_2_O_2_ production during re-oxygenation, while metabolic NADP reduction is efficiently restored. The reduction of the NADP pool suggests that the electron influx into the NADP pool by metabolism more than compensates for the electron efflux to the anti-oxidant systems, and that glutathione reductase activity, catalysing the reduction of glutathione from NADPH, is limiting under the specific conditions of re-oxygenation after hypoxia.

We reasoned that the diametrically opposed redox responses of the NADP and glutathione pools to reoxygenation, i.e. NADP reduction and glutathione oxidation, represented an opportunity to benchmark the recently introduced biosensor NERNST against NAPstars and roGFP2-Grx1 probes *in vivo*. NERNST is based on a redox-sensitive GFP (roGFP2), which is known to respond strongly to changes in glutathione redox state (*E*_GSH_) in a reaction efficiently catalysed by endogenous glutaredoxins in most cell compartments, including the plant cytosol [19–22]. This raises the crucial question of whether NERNST really reports cytosolic NADP redox state *in vivo* or whether its specificity is compromised by equilibration with *E*_GSH_ as catalysed by the endogenous glutaredoxins (Grx). We thus repeated the hypoxia-reoxygenation experiments in leaves with NAPstar4.3, NAPstarC, Peredox, roGFP2-Grx1 and NERNST sensors (**Figure S10**) The NERNST response was almost indistinguishable from a roGFP2-Grx1 response (**Figure S10C,D**) consistent with NERNST responding predominantly to changes in *E*_GSH_ in the Arabidopsis cytosol. Since high Grx activity, through several different Grx isoforms, is common in the cytosol of plants in general, we conclude that NERNST is not an NADP redox state sensor in plants and functions primarily as a glutathione redox state reporter.

### The glutathione pathway mediates high capacity NADPH-dependent anti-oxidative electron flux

Having established the functionality of NAPstars in yeast, mammalian cells and plants, we next sought to address a longstanding question in redox biology, namely, the role of glutathione in cellular anti-oxidative responses. Even though glutathione is often loosely referred to as a key cellular anti-oxidant, its mechanistic roles within cells have been remarkably difficult to pin down and a matter of a longstanding debate. Importantly, the thioredoxin pathway has been considered to be the primary reductive system, with glutathione seen as having a back-up or auxiliary role [38–40].

Upon exposure to exogenous oxidants, such as H_2_O_2_ or diamide, there is a heightened demand for NADPH. Both the thioredoxin reductase/thioredoxin and glutathione reductase/glutathione/glutaredoxin pathways (ascorbate-glutathione cycle in plants) [41] use electrons from NADPH to reduce cellular disulfide bonds and certain reactive oxygen and reactive nitrogen species. Nonetheless, monitoring the relative flux of electrons through these two pathways *in vivo* under pro-oxidative conditions has proven challenging. To address the question about the relative contribution to anti-oxidant electron flux we made use of genetic or chemical inhibition of the glutathione and thioredoxin-dependent pathways and monitored the cytosolic NADP redox dynamics in response to exogenous oxidants in living cells and tissues.

First, we examined yeast cells. We monitored the response of NAPstarC, NAPstar3 and PeredoxDS probes expressed in wild-type cells, in cells deleted for the *GLR1* gene (encoding glutathione reductase), and in cells deleted for *TRX1* and *TRX2* (encoding the two cytosolic thioredoxins). The probe responses in all cells were monitored in response to the addition of exogenous diamide at concentrations ranging from 0–5 mM (**Figure 5A–E**). Diamide was chosen instead of H_2_O_2_ due to the extremely small deflections in cytosolic NADPH redox homeostasis detected in yeast in response to peroxides (**Figure 2**). In wild-type cells, we observed a strong oxidation of the cytosolic NADP pool upon treatment with 2 and 5 mM diamide (**Figure 5B**). Intriguingly however, in Δ*glr1* cells, we observed almost no detectable NADP oxidation at all, except for a slight response of the NAPstar3 probe to the highest diamide concentration (5 mM) (**Figure 5C,E**). Deletion of both cytosolic thioredoxins also led to less severe NADP oxidation following diamide treatment in comparison to that observed in wild-type cells (**Figure 5D,E**). However, the amplitude change of NAPstar3 TS/mC ratio was much larger than observed in Δ*glr1* cells (**Figure 5E**). The response of PeredoxDS was similar in wild-type, Δ*glr1* and Δ*trx1*Δ*trx2* cells, whilst NAPstarC showed no response (**Supplementary Fig 11**). Our results indicate that genetic impairment of both the thioredoxin and glutathione-dependent pathways can hinder the consumption of NADPH upon pro-oxidative challenges in the yeast cytosol, but the impact of blocking the glutathione pathway is more pronounced.

**Figure 5.**
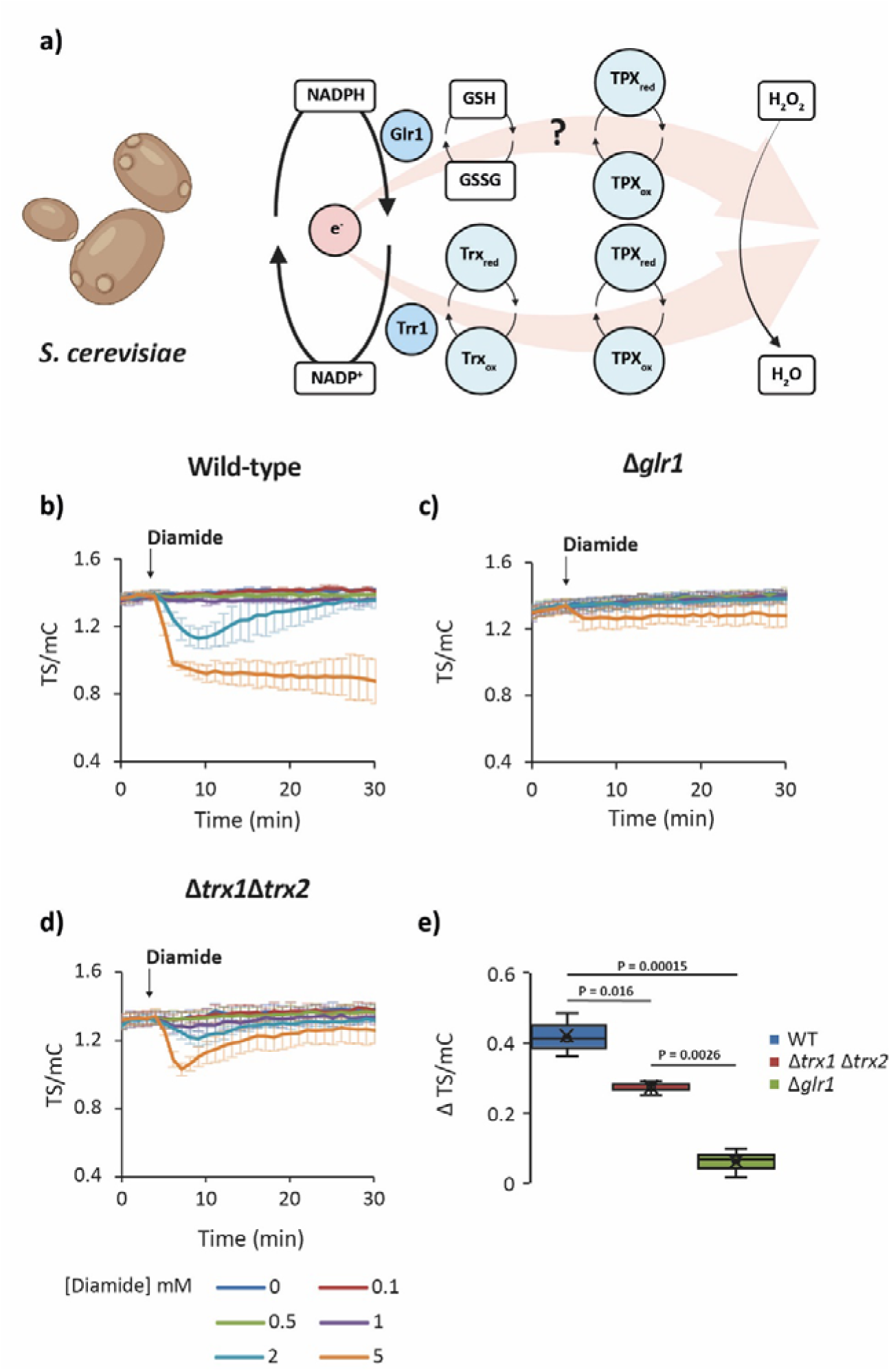
Glutathione reductase deletion protects against diamide-induced NADP oxidation in yeast. **a,** Cartoon showing two possible pathways by which electrons from NADPH can be transferred to H_2_O_2_ in the yeast cytosol, TRX = thioredoxin, TPX = thiol peroxidase, GR = glutathione reductase. Created with BioRender.com. **b–d,** Response of a cytosolic NAPstar3 probe to exogenous diamide at the indicated concentrations in wild-type (**b**), Δ*glr1* (**c**), and Δ*trx1*Δ*trx2* (**d**) cells (n=3 measurements made with cells derived from independent cultures). **e,** Box and whisker plot showing the change in TS/mC before and after treatment with 5 mM diamide. P-values are derived from a one-sided ANOVA test. In all panels, data are presented as mean ± s.d.

We again saw a chance to test the response of the NERNST sensor. Lack of coupling between the NADP redox status and *E*_GSH_ in Δ*glr1* cells, provides a powerful genetic system to separate NADP and *E*_GSH_-dependent responses. At acute oxidative challenge, *E*_GSH_ is expected to respond more strongly (and recover less quickly) in Δ*glr1* cells, whilst NADP redox status is expected to respond less strongly (and recover more quickly). NERNST responses to both H_2_O_2_ and diamide were strongly increased in Δ*glr1* cells in comparison to wild-type cells, suggesting that *E*_GSH_ dominates the NERNST response in the cytosol of living yeast cells. Consistently, the NERNST response was very similar to roGFP2-Grx1 responses **(Figure S12 and 13**). These results show in a second biological system tested that *in vivo* NERNST responses are usually dominated by the catalysed interaction of its roGFP2 domain with the glutathione redox couple. The slightly slower kinetics of NERNST oxidation compared to the Grx1-roGFP2 probe likely stem from NERNST relying on endogenous glutaredoxins in the yeast cytosol to catalyse the equilibration of its roGFP2 domain with the glutathione redox couple. In Grx1-roGFP2, the genetic fusion of a glutaredoxin to roGFP2 increases the effective local glutaredoxin concentration by ∼1000-fold [42], thereby allowing roGFP2 to equilibrate more rapidly with changes in *E*_GSH_.

We next asked whether the different impacts of impairing the glutathione and thioredoxin pathways are specific to yeast or also hold true in other eukaryotic systems. Therefore, we monitored the response of NAPstar4.3 in *A. thaliana* wild-type plants, as well as in plants lacking the two cytosolic NADPH-dependent thioredoxin reductases *ntr a/b* or cytosolic glutathione reductase *gr1* (**Figure 6 and Figure S14**). We observed an H_2_O_2_ concentration-dependent NADPH consumption in leaf discs from wild-type plants, which was similar in *ntra/b* plants, but completely absent in *gr1* plants. A dominant role of the glutathione system over the thioredoxin system in mediating anti-oxidant electron flux *in planta* is consistent with our observations in yeast, and the concept of the ascorbate-glutathione cycle as the dominant route of NADP-dependent anti-oxidant electron flux in plants.

**Figure 6.**
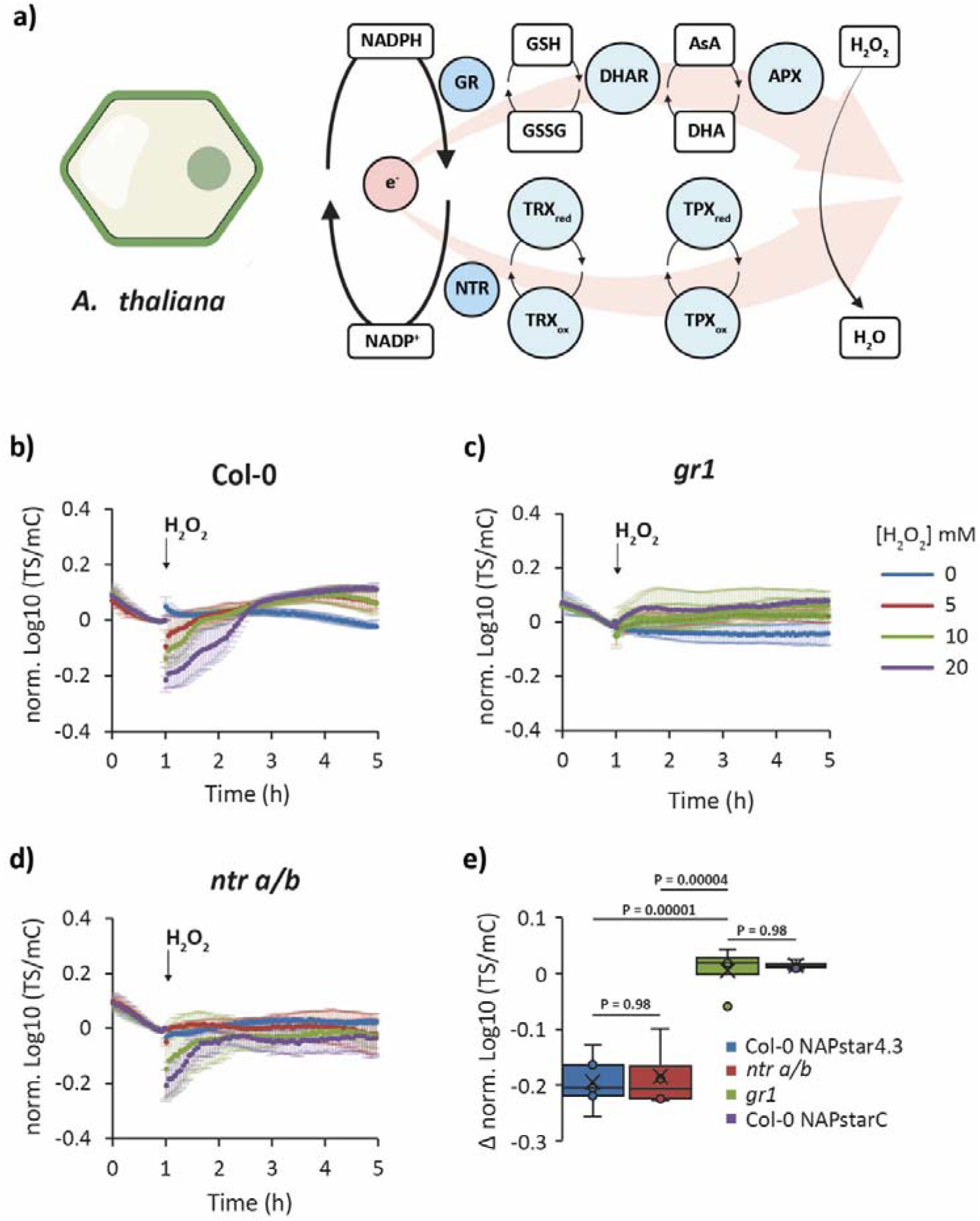
Glutathione reductase deletion protects against H_2_O_2_-induced NADP oxidation in plants. **a,** Cartoon showing two possible pathways by which electrons from NADPH can be transferred to H_2_O_2_ in the plant cytosol, TRX = thioredoxin, TPX = thiol peroxidase, GR = glutathione reductase, DHAR = dihydroascorbate reductase, APX = ascorbate peroxidase. Created with BioRender.com. **b–d,** Response of a cytosolic NAPstar4.3 probe to exogenous H_2_O_2_ at the indicated concentrations in wild-type (Col-0) plants (**b**), *gr1* (**c**), *ntr a/b* (**d**) plants, and the control construct NAPstarC in wild-type plants (**e**) (n=4–5 leaf discs from individual plants). **f,** Box and whisker plot showing the change in TS/mC before and after treatment with 20 mM H_2_O_2_. P-values are derived from a one-sided ANOVA test. In all panels, data are presented as mean ± s.d.

Finally, we asked whether this principle may be generalized and tested the impact of glutathione reductase deletion on NADP redox responses in mammalian cells lines. (**Figure 7 and Figure S15**). To this end, we monitored the response of NAPstar3b in HEK293 cells and in HEK293 cells with a CRISPR-Cas9-mediated disruption of GSR (encoding glutathione reductase). The response of NAPstar3b in several hundred individual cells was imaged using a plate-reader-based fluorescence microscopy setup. Cells were treated with diamide at 100 or 500 µM (**Figure 7B–D**) and with H_2_O_2_ at 50 or 100 µM (**Figure 7E–G**). Consistent with our observations in yeast and plants, we observed cytosolic NADP oxidation in control cells in response to both H_2_O_2_ and diamide. However, we observed no detectable NAPstar3b response to either oxidant in GSR KO cells. Finally, we tested the impact of chemical inhibition of glutathione reductase and thioredoxin reductase using 1,3-bis-(2-chlorethyl)-l-nitroso-urea (BCNU) and auranofin, respectively (**Figure 7 H–J**). In HeLa cells treated with auranofin for 60 min prior to 100 µM H_2_O_2_ addition, we observed no difference in NAPstar3b response compared to untreated control cells. In contrast, in cells pre-treated for 60 min with BCNU prior to H_2_O_2_ addition, we observed no NAPstar3b response at all, consistent with our observations of genetic disruption of glutathione reductase in yeast, plants, and HEK293 cells. In summary, we conclude that the glutathione system is the dominant mediator of anti-oxidative electron flux in pro-oxidant conditions across different eukaryotic kingdoms.

**Figure 7.**
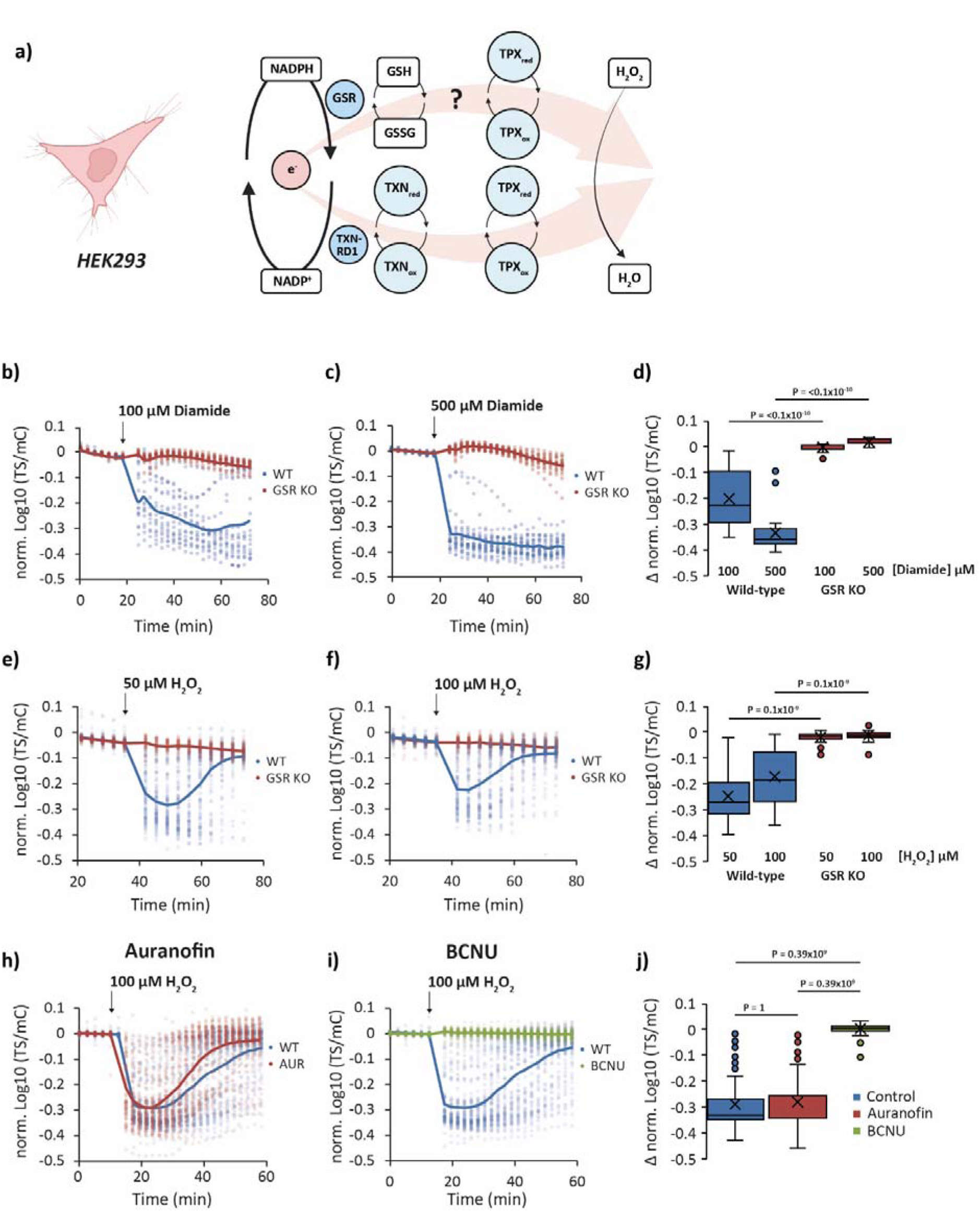
Glutathione reductase deletion protects against H_2_O_2_-induced NADP oxidation in HEK293 cells. **a,** Cartoon showing two possible pathways by which electrons from NADPH can be transferred to H_2_O_2_ in the mammalian cell cytosol, TXNRD1 = thioredoxin reductase, TRX = thioredoxin, TPX = thiol peroxidase, GSR = glutathione reductase. Created with BioRender.com. **b,c,** Response of NAPstar3b expressed in the cytosol of wild-type and glutathione reductase deleted (GSR KO) cells to 100 µM diamide (**b**) (n=17 or 18 individual cells for wild-type and GSR KO, respectively, measured in two separate experimental repeats) or 500 µM diamide (**c**) (n=22 or 19 individual cells for wild-type and GSR KO, respectively, measured in two separate experimental repeats. **d,** Box and whisker plot showing the change in TS/mC before and after treatments in **b.** and **c. e,f,** Response of NAPstar3b expressed in the cytosol of wild-type and glutathione reductase deleted (GSR KO) cells to 50 µM H_2_O_2_ (**e**) (n=103 or 46 individual cells for wild-type and GSR KO respectively, measured in four and three separate experimental repeats) or 100 µM H_2_O_2_ (**f**) (n=98 or 76 individual cells for wild-type and GSR KO, respectively, measured in four and five separate experimental repeats. **g,** Box and whisker plot showing the change in TS/mC before and after treatments in **e.** and **f. h,i,** Response of NAPstar3b in the cytosol of HEK293 cells, treated either with a vehicle control, the thioredoxin reductase inhibitor auranofin (**h**) or the glutathione reductase inhibitor 1,3-bis(2-chloroethyl)-1-nitrosourea (BCNU) (**i**) (n= 85, 97, 79 individual cells for control, auranofin and BCNU treatments respectively, monitored in the course of 3 experimental repeats). **j,** Box and whisker plot showing the change in TS/mC before and after treatments in **h.** and **i.**

## Discussion

Here we report on the development of the NAPstar family of genetically encoded fluorescent probes for in cell monitoring of the NADP redox state. NAPstars are functional in yeast, plants and mammalian cells. Importantly, unlike other currently available NADP sensors, including iNaps and Apollo-NADP^+^ [15, 16], NAPstars respond to changes in both NADPH and NADP^+^ and are *bona fide* probes for the NADP redox state (**Figure 1; Figure S1-4**). NAPstars exhibit several other advantages over currently available NADP sensors including pH-resistance (**Figure S3**), which affords the possibility to monitor more reliably in situations where pH can respond readily to environmental changes, for example in the yeast cytosol [43] or in the plant cytosol during hypoxia or changes in illumination [36, 44]. Furthermore, NAPstars contain both of the Rex domains necessary for NADPH or NADP^+^ binding within one probe molecule. NAPstars are thus self-contained probes and unlike iNap sensors do not rely on probe dimerization to function [15]. This greatly increases the possibility for engineering of probe properties such as NADPH or NADP^+^ binding affinity and nucleotide specificity by creating NAPstar probes composed of two differently mutated Rex domains. The mixed Rex domain NAPstar4.3 for example has no detectable binding affinity for NADH and has a *K*_dNADPH_ ≈ 43 µM and a *K*_r(NADPH/NADP+)_ ≈ 0.28, which is approximately one order of magnitude lower than other NAPstar sensors (**Figure 1**) and allows fully dynamic measurement of NADP oxidation and reduction in the yeast cytosol during the yeast metabolic cycle and in the plant cytosol during illumination and hypoxia (**Figure 3 and 4**). The self-contained nature of NAPstars also allows them to function independently of their expression level whereas probes that rely on dimerization may potentially have issues at lower expression level. We also demonstrate that NAPstars are amenable to FLIM measurement, with a large change in fluorescence lifetime between the NADPH and NADP^+^-bound states, as exemplified by NAPstar4.3 (**Figure S5**). Finally, in common with other NAD(P) probes, but unlike genetically encoded probes for redox species such as H_2_O_2_ [6, 25, 31, 45] or glutathione [19], NAPstars *bind* their target redox species fully reversibly but do not *react* with it. *In vivo*, NAPstars are one amongst many other endogenous NADP-binding proteins decreasing the likelihood that NAPstars perturb the NADP pool to any considerable extent.

NERNST is a recently published sensor for measuring the NADP redox state [18]. NERNST is based on a genetic fusion between redox-sensitive green fluorescent protein 2 (roGFP2) [18] and NADPH-thioredoxin reductase C (NTRC) from rice (*Oryza sativa*). Whilst an elegant concept, there are legitimate reasons to question the specificity of NERNST in many biological contexts. RoGFP2 has been shown to rapidly equilibrate with the glutathione redox couple in a reaction catalysed by endogenous redox-active glutaredoxins [19, 20, 22]. An interaction with glutathione/glutaredoxin has been observed in all roGFP-based sensors generated to date [19, 31, 45–49] and it is unclear why this interaction should not also occur in the context of the NERNST probe. Here, we employed two different experimental settings in which glutathione and NADP redox responses are diametrically opposed to test whether NERNST responds predominantly to NADP or glutathione redox changes. In both experiments, i.e. the differential response of the glutathione and NADP redox states to oxidative challenge in Δ*glr1* and wild-type yeast and during re-oxygenation in plants, the behaviour of NERNST closely mirrored that of the well-established *E*_GSH_ sensor, roGFP2-Grx1, and was opposite to that of the NAPstars. We thus conclude that NERNST predominantly responds to changes in *E*_GSH_ *in vivo*. Therefore NERNST is not an NADP redox state sensor the cellular contexts assessed here, and this conclusion can be plausibly extrapolated to most cellular systems where endogenous cytosolic Grx activity is typically high.

As with every methodological approach, NAPstars also have limitations of which it is important the user is aware. As with any genetically encoded probe, NAPstars require genetic manipulation of the organism in which the probe is intended to be used. NAPstars are relatively large constructs, which may limit their targetability or usability in some subcellular locations, such as experienced for targeting Peredox to the mitochondrial matrix. Finally, it is important to be aware that NAPstars monitor the soluble pool of NADPH and NADP^+^, they do not ‘see’ protein-bound NADP, which may constitute a significant fraction of total NADP in some situations [32].

The cytosolic NADP redox state is typically considered to be highly reduced, with NADPH/NADP^+^ ratios of 50–100:1 being standard textbook values [50, 51]. However, a very wide-range of values for cytosolic NADPH/NADP^+^ has been reported in the literature [32, 52–54]. This raises questions regarding what the actual cytosolic NADP redox state is and/or how variable it might be under different conditions or in different organisms. NAPstars have *K*_r(NADPH/NADP+)_ values ranging from 0.0006–0.28. This means that they can respond to changes in NADPH/NADP^+^ ranging from about 1:1000 to about 5:1. The fact that NAPstars allow for fully dynamic measurements *in vivo*, would support the conclusion that the cytosolic NADP redox state lies somewhere within this range and thus is much more oxidised than often considered. As NAPstar4.3 was best suited for fully dynamic measurements in plants leaves, and in glucose-limited continuous yeast cultures, we can conclude that cytosolic NADPH/NADP^+^ in these systems is in the range of about 1:10 to about 5:1, which would give cytosolic *E*_NADP_ ranging from −290 mV to –340 mV. This is intriguing as numerous measurements of the cytosolic *E*_GSH_ report similar values, typically in the range of −300 mV to –320 mV [55]. The apparent overlap in redox potentials could suggest that the cytosolic glutathione and NADP redox couples are able to rapidly equilibrate with each other, for example by the action of glutathione reductase. Supporting this hypothesis, we observed that despite the more oxidised than expected cytosolic NADP redox state, it seems to be coupled with a remarkable robustness to perturbation by exogenous oxidants in both yeast and mammalian cells. This is strikingly similar to our previous observations on the robustness of cytosolic glutathione redox homeostasis [21, 56] and raises the exciting possibility that glutathione reductase may facilitate the rapid bi-directional interaction between the glutathione and NADP redox couples, which could suggest that the highly abundant glutathione pool might effectively serve as a ‘reservoir’ of available electrons to supply the NADP pool under conditions of extreme oxidative challenge. It will be fascinating to explore the *in vivo* crosstalk of those key redox couples further in the future, and to understand situations in which the glutathione and NADP redox couples are clearly not coupled, including those documented in this manuscript.

In the yeast and mammalian cytosol, the thioredoxin pathway was considered to be the major reductive pathway with the glutaredoxin/glutathione system having only a back-up or auxiliary role [38, 40]. In the plant cytosol there has been evidence for a major anti-oxidant electron flux through the glutathione system, specifically via the ascorbate-glutathione cycle [41, 57]. Yet, it was proposed that the only essential function of glutathione is in Fe-S cluster biogenesis [38]. Nonetheless, only micromolar amounts of glutathione are required for Fe-S cluster biogenesis and the biological reason for the millimolar concentrations of glutathione present in cells across eukaryotic kingdoms remains unclear [58]. Furthermore, indirect evidence from several studies supports an important role of glutathione as a reductant, especially in response to acute oxidant challenges. For example, glutathione disulfide (GSSG) is readily produced in cells in response to exogenous oxidants, supporting the concept that GSH is acting as a reductant in these situations [21, 57]. Cellular GSSG accumulation is limited by the fact that cells employ multiple redundant reductive pathways to reduce cytosolic GSSG or to actively export GSSG to the extracellular environment, for example by MRP1 in mammalian cells or to alternative subcellular compartments, for example by Ycf1 in yeast [21, 59]. Moreover, GSH has been shown to be able to act as a source of reductive power for typical 2-Cys peroxiredoxins, including human PRDX2 [60] and Arabidopsis PRXIIB, C and D [61]. The relative importance of the glutathione system compared to the more important thioredoxin system may be further amplified under acute oxidative challenge as thioredoxins or thioredoxin reductases may readily become limiting and/or oxidised and thus inactive [62], which would strongly limit the capacity of the thioredoxin pathway to act as a conduit of anti-oxidative electron flux.

Here we used NAPstar sensors, in yeast, mammalian cells, and plants in combination with genetic and chemical inhibition to re-visit the question of the relative importance of the glutathione and thioredoxin systems as mediators of anti-oxidative electron flux during acute oxidative challenges. In yeast and plant, we observed almost no NADPH consumption upon oxidative challenge with H_2_O_2_ or diamide in the absence of a cytosolic glutathione reductase (**Figure 5 and 6**). Likewise, in mammalian cells, deletion of glutathione reductase or chemical inhibition with BCNU, led to almost complete loss of NADPH consumption with both H_2_O_2_ and diamide (**Figure 7**). In contrast, genetic or chemical ablation of the thioredoxin pathway, had a much smaller effect in all three organisms. We interpret these results to mean that the glutathione system acts throughout the eukaryotic kingdom as the major mediator of anti-oxidative electron flux in response to acute oxidative challenge. It will be exciting in the future to utilise NAPstar sensors to investigate the relative importance of the glutathione and thioredoxins pathways under situations of endogenously generated oxidative challenge, such as frequently occurring stress situations.

In conclusion, we have developed and extensively characterised the NAPstar family, which are the first *bona fide* biosensors of the NADP redox state. NAPstar are bright, pH-resistant and specific NADP redox state sensors, which can be measured either by standard fluorescence spectroscopy-based approaches or by FLIM. We have used NAPstars to uncover new biology in yeast, plant and mammalians systems, including a conserved and unexpected dominance of the glutathione system as a mediator of anti-oxidative electron flux. We are confident that NAPstars will drive discovery and understanding of a broad range of novel NADP biology and its synthetic re-wiring in the future.

## Acknowledgements

B.M. and L.P.R. gratefully acknowledge funding in the context of the Saarland University NanoBioMed Method Development Seed Funding. The Deutsche Forschungsgemeinschaft (DFG, German Research Foundation) provided funds for research in the lab of Bruce Morgan through the grant MO 2774/6-1 project number 505680640. The DFG provided funds for research in the Laboratory of Jan Riemer through the grants RI2150/5-1 project number 435235019, RI2150/2-2 project number 251546152, RTG2550/1 project number 411422114, and CRC1218 project number 269925409. Markus Schwarzländer also acknowledges the DFG for funding through the infrastructure grant INST211/903-1 FUGG, and the grants SCHW1719/9-1 project number 508398975, SCHW1719/10-1 project number 507704013 and SCHW1719/11-1 project number 507704013. Joris Messens is funded with a VIB grant. We thank Andreas J. Meyer, Stephan Wagner and José Manuel Ugalde (Bonn) for stimulating discussions about NERNST specificity.

## Materials and Methods

### Plasmid construction

Amino acids mutations to switch Rex-domain specificity from NAD to NADP were selected based on a previous screen [15]. All constructs and gene sequences used in this study were either commercially synthesized, generated by standard molecular cloning approaches or were generated in previous studies (**Supplementary Information**; **Supplementary Table 1–5**). Primer sequences are provided in **Supplementary Table 6**. All sequences were confirmed by commercial sequencing (Eurofins Genomics, Ebersberg, Germany).

#### Cloning and site-directed mutagenesis

NAPstar sequences for plant expression were commercially synthesized with codons optimized for plant expression (GenScript Biotech, Rijswijk, Netherlands) and inserted into pDONR207 (LIFE Technologies, Carlsbad, CA, United States) (**Supplementary Table 1**). The NAPstar3b variant was generated by site-directed mutagenesis on the pDONR207 NAPstar3 plasmid using the primer pair Pr1/Pr2 (**Supplementary Table 6**). For plant expression all constructs were subcloned into pSS02 (derivative of pMDC32; [23, 63]) using gateway cloning (**Supplementary Table 2**). Plant NAPstar sequences exhibit a pre-existing *HindIII* restriction site within the cpT-Sapphire (TS) domain. Using this restriction site, together with an *ApaI* restriction site in the backbone of pDONR207, the N-terminal Rex domain of NAPstar4 was fused to the second, C-terminal, Rex domain of NAPstar3 to generate NAPstar4.3. For bacterial expression, all plant NAPstar sequences were subcloned by gateway cloning [64] into pETG10a (Invitrogen, Carlsbad, CA, United States) (**Supplementary Table 3**).

All coding sequences for NAPstars, Peredox, HyPer7 and NERNST were codon optimized for expression in *Saccharomyces cerevisiae*, synthesized and delivered in a pUC57 plasmid (GenScript Biotech, Rijswijk, Netherlands). For yeast expression, all coding sequences were subcloned into an empty p413TEF plasmid [65] using *XbaI* and *XhoI* restriction sites (**Supplementary Table 5**). A pre-existing *ClaI* restriction site in the TS sequence was utilised to fuse the first, N-terminal, Rex domain of NAPstar4 to the second, C-terminal, Rex domain of NAPstar3 to generate the NAPstar4.3 construct. The p413TEF roGFP2-Grx1 plasmid was generated by subcloning from a p415TEF roGFP2-Grx1 plasmid. The PeredoxDS construct was obtained after two rounds of mutagenesis on the p413TEF Peredox plasmid using primer pairs Pr3/Pr4 and Pr5/Pr6 (**Supplementary Table 6**). Site-directed mutagenesis was performed using a standard PCR-based protocol with S7 Fusion Polymerase (Biozym Scientific GmbH, Hessisch Oldendor, Germany). Methylated template DNA was digested using *DpnI* (NEB, Ipswich, MA, United States), before heat-shock transformation into chemically competent *E. coli* Top10 cells.

Codon optimized sequences of NAPstar3b and HyPer7 for expression in mammalian cells were commercially synthesized (**Supplementary Table 4**) (GenScript Biotech, Rijswijk, Netherlands) and delivered in pcDNA3.1(+) plasmids.

### AlphaFold2 structure prediction of monomeric NapStar

The structural prediction was performed with AlphaFold2 with ColabFold. The following script was used for running the structural prediction process: colabfold_batch --model-type AlphaFold2--num-recycle 48 --amber --use-gpu-relax) [66] using the amino acid sequence of NAPstar3 as input (**Supplementary List 1**). In the prediction process, 48 recycling steps were employed, serving as iterations where the model fine-tuned its predictions to enhance accuracy. The refinement stage utilized AMBER (Assisted Model Building with Energy Refinement), a force field commonly used in molecular dynamics simulations. Additionally, a relaxation process was implemented, optimizing the predicted structures further to attain more realistic and energetically favorable conformations. Visualisation was performed using PyMol software (https://pymol.org).

### Expression and purification of recombinant proteins

Protein expression and purification were performed as previously described [23, 67] with the following modifications. Proteins were isolated from either an *E. coli* BL21 (DE3) ArcticExpress strain or from an *E. coli* Rosetta 2 (DE3) strain expressing pETG10a NAPstar or pETG10a Peredox. Isopropyl β-D-1-thiogalaytopyranoside (IPTG) was added at a concentration of 0.5 mM for overnight induction at 20°C. Bacterial pellets were obtained by centrifugation for 15 min at 3,220 *g* and pellets were resuspended in 50 mM Tris-HCl, 100 mM NaCl, 0.5 mM MgCl_2_ (pH 7.5). Cell lysates were centrifuged at 12,100 *g* for 15 min at 4°C. Clarified lysates were incubated with Ni-NTA beads. Beads were washed with 20 mM and 40 mM imidazole steps and protein was eluted using 250 mM imidazole in 50 mM Tris-HCl, 100 mM NaCl, 0.5 mM MgCl_2_ (pH 7.5).

### *In vitro* characterization of NAPstar sensor

*In vitro* assays were performed in 50 mM Tris-HCL, 100 mM NaCl, 0.5 mM MgCl_2_ at pH 7.5, except for pH titration and enzyme-coupled assays experiments. For all *in vitro* assays the NAPstar sensor protein was quantified by Bradford assay [68] and the sensor protein was used at a final concentration 240–480 nM.

*In vitro* assays were performed using NUNC 96-well plates (VWR International GMBH, Darmstadt, Germany) and a CLARIOstar plate reader (BMG Labtech, Ortenberg, Germany). Emission spectra were measured in 2 nm steps between 425 nm and 601 nm for TS and between 570 nm and 650 nm for mCherry, after excitation with 400 ±5 nm and 540 ±10 nm, respectively. The excitation spectra were measured in 2 nm steps between 350 nm and 494 nm for TS and between 400 nm and 588 nm for mCherry at a fixed emission detection at 520 ±5 nm and 615 ±9 nm, respectively. For measurements over time the NAPstar biosensors were excited at 400 ±5 nm and 540 ±10 nm with detection of the emission at520 ±5 nm and 615 ±9 nm for TS and mCherry respectively. NAD(P)H autofluorescence was measured with excitation at 340 ±7.5 nm and the emission was detected at 445 ±10 nm.

The pH titration experiments were performed in 100 mM HEPES, 150 mM NaCl_2_, 0.5 mM MgCl_2_ and pH between 6 and 9 adjusted by KOH, with the actual pH measured post-hoc. For enzyme-coupled assays, 50 µU alkaline phosphatase FastAP (EF0651, Thermofisher Scientific, Waltham, Massachusetts, United States), 5–15 µU glutathione reductase (G3664-100UN, Sigma Aldrich, St. Louis, MO, United States)) and 1.07 ng.µL^-1^ isocitrate dehydrogenase (I2411-10UG, Sigma Aldrich, St. Louis, MO, United States)) were used. The alkaline phosphatase assay was performed in 10 mM Tris-HCl, 5 mM MgCl_2_, 100 mM KCl, 0.02% (v/v) Triton-X-100 and 0.1 µg.µL^-1^ BSA, pH 8.0. The glutathione reductase assay was performed in 50 mM Tris-HCl, 100 mM NaCl, 0.5 mM MgCl_2_ and 1 mM EDTA, pH 7.5. The isocitrate dehydrogenase assay was performed in 50 mM Tris-HCl, 100 mM NaCl, 5 mM MgCl_2_, 2.4 mM KCl and 0.65 µg/µl BSA, pH 8.0.

### Growth of yeast strains

The *S. cerevisiae* BY4742 (MATα *his3*Δ1 *leu2*Δ0 *lys2*Δ0 *ura3*Δ0) strain background [69] was used for all fluorescence plate-reader experiments. For all fermentor experiments a CEN.PK113-1A yeast strain was used [30]. All yeast strains used in this study (**Supplementary Table 7**) were grown as previously described in Hartwell’s Complete (HC) medium containing 2% (w/v) glucose as carbon source [70] and lacking histidine for plasmid selection unless stated otherwise.

### Construction of yeast strains and transformation

Yeast gene deletions were generated by PCR-based homologous recombination [71] (**Supplementary Table 7**). An antibiotic resistance cassette was PCR-amplified from either pFA6α *natNT2* or pFA6α *kanMX4* plasmids using primers designed to have 40–50 base-pairs of homology directly up- and down-stream of the gene to be deleted. Deletion of *ZWF1, TRX1* and *TRX2* in a BY4742 background was achieved using the primer pairs Pr7/Pr8, Pr9/Pr10, Pr11/Pr12 respectively. Deletion of *HIS3* in a CEN.PK113-1A background was achieved by using the primer pair Pr13/Pr14 (**Supplementary Table 6**).

The PCR products of these reaction were transformed into yeast cells using a standard lithium acetate-based approach. Briefly, cells were grown in YPD until logarithmic phase, harvested by centrifugation at 1,000 *g* for 3 min at room temperature and resuspended in 200 µl ‘One-step-transformation’ buffer containing 40% polyethylene glycol 3,350 (Sigma Aldrich, St. Louis, MO, United States), 0.2 M lithium acetate (Sigma Aldrich, St. Louis, MO, United States) and 0.1 M dithiothreitol (DTT; AppliChem GmbH, Darmstadt, Germany). Following the addition of 10 µl salmon testes single stranded DNA and the PCR product, cells were incubated with continuous shaking for 30 min at 45°C. Subsequently, cells are transferred into fresh YPD and grown overnight before plating onto YPD plates containing the appropriate antibiotic. After two rounds of selection, gene deletions were confirmed by PCR on genomic DNA using primers binding ∼400 base pairs up- and down-stream of the gene of interest. DNA was extracted by heating yeast cells at 96°C in 0.2% (w/v) SDS for 10 min. Subsequently cells were vortexed thoroughly and spun down at 11,000 *g* for 1 min. 1 µL of genomic DNA-containing supernatant was used as template in the PCR reaction.

### Plate reader-based fluorescence measurements in *S. cerevisiae*

Yeast strains were either transformed with empty p413TEF plasmids for fluorescence background subtraction or p413TEF plasmids containing the indicated genetically-encoded sensor. For plasmid transformation into yeast cells, a standard lithium acetate-based approach was used as described above. Yeast cells are harvested and resuspended in 100 µl ‘One-step-transformation’ buffer before 5 µl of salmon testes single stranded DNA and ∼200 ng of Plasmid DNA was added. Cells were then incubated with continuous shaking at 45°C for 30 min, before plating on HC medium containing 2% (w/v) glucose as carbon source lacking histidine to ensure plasmid retention. Plates were then incubated for 2 days at 30°C.

Fluorescence of NAPstar and Peredox constructs was measured using excitation at 399 ±10 and 510 ±10 nm with emission at 578 ±15 and 619 ±15 nm for the TS and mCherry domains respectively. For data calculation background fluorescence was subtracted before intensiometric TS fluorescence signals were divided by mCherry signals to allow for a ratiometric, probe expression-independent readout. For all roGFP2-Grx1 and NERNST measurements, roGFP2 was excited at either 400 ±15 nm or 480 ±15 with emission at 520 ±20 nm. Calculation of the degree of oxidation (OxD roGFP2) was performed according to equation 1:

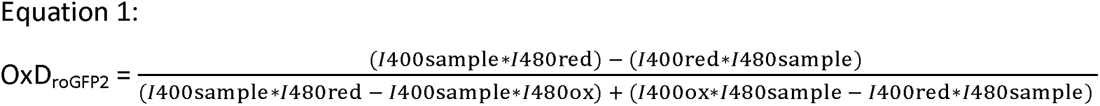

For plate-reader experiments, yeast cultures were grown to late logarithmic phase (OD_600_ = 3–4) in liquid HC medium lacking histidine for plasmid retention. Cells were then harvested by centrifugation at 1,000 *g* for 3 min at room temperature and resuspended in 100 mM MES/Tris buffer, pH 6, to an OD_600_ of 7.5. 200 µL aliquots were then transferred to the wells of a NUNC flat-bottomed 96-well microtiter plate (VWR International GMBH, Darmstadt, Germany). Subsequently, the plate was centrifuged at 30 *g* for 5 min at room temperature, so that cells form loose pellets at the bottom of each well. The measurement was initiated by addition of the indicated treatment and fluorescence changes of each probe were monitored for 60 min using a CLARIOstar (BMG Labtech, Ortenberg, Germany) fluorescence plate-reader. All experiments were performed at least 3 times with cells from independent cultures.

### Online monitoring of metabolite dynamics in continuous fermentor cultures

A Biostat A fermentor (Sartorius Stedim Systems GmbH, Guxhagen, Germany) was used to measure population-synchronized metabolic oscillations in continuous cultures with respect to the yeast metabolic cycle [30]. For all fermentor-based experiments a CEN.PK113-1A strain background was used. A CEN.PK113-1A Δ*his3* strain was generated to allow for retention of *HIS3*-containing p413TEF plasmids. This strain was transformed with p413TEF plasmids with the indicated genetically encoded probes. Culture media consisted of 10 g.L^-1^ glucose, 1 g.L^-1^ yeast extract, 5 g.L^-1^ (NH_4_)_2_SO_4_, 2 g.L^-1^ KH_2_PO_4_, 0.5 g.L^-1^ MgSO_4_.7H_2_O, 0.1 g.L^-1^ CaCl_2_.2H_2_O, 0.02 g.L^-1^ FeSO_4_.7H_2_O, 0.01 g.L^-1^ ZnSO_4_.7H_2_O, 0.005 g.L^-1^ CuSO_4_.5H_2_O, 0.001 g.L^-1^ MnCl_2_.4H_2_O, 2.5 mL 70% H_2_SO_4_ and 0.5% (v/v) Antifoam 204 (Sigma Aldrich, St. Louis, MO, United States). Fermentor runs were initiated by the addition of a 20 ml starter culture grown at 30°C to stationary phase in HC medium lacking histidine for plasmid retention. The fermentor was run with a working volume of 800 ml at 30°C with constant aeration of 1 L.min^-1^ and stirring at 530 rpm. The automated addition of 10% (w/v) NaOH maintained a constant pH of 3.4. The culture was initially run in batch-culture mode until ∼6 h after the exhaustion of the carbon source as determined by continuous and automated monitoring of the oxygen saturation within the culture vessel every 10 s. A continuous culture was subsequently initiated by addition of fresh medium at a dilution rate of 0.05 h^-1^. To enable the continuous ‘online’ monitoring of genetically-encoded probes, an in-house coupled fermentor-fluorimeter system was previously developed [30]. A peristaltic pump was used to continuously pump culture from the fermentor through a flow cell (Type 71-F, Starna GmbH, Pfungstadt, Germany), which was inserted into a JASCO FP-6500 spectrofluorimeter (JASCO, Oklahoma City, OK, United States) before media returned to the fermentor vessel. Fluorescence of PeredoxDS and NAPstar probes was measured at fixed excitation and emission wavelengths at 399 ±10 and 510 ±10 nm or 578 ±10 and 619 ±10 nm for the TS and mCherry domains respectively. Fluorescence of HyPer7 was measured at fixed excitation wavelengths of 400 ±10 nm and 490 ±10 nm with emission monitored at 520 ±10 nm. A slit width of 10 nm was used, and fluorescence measurements were continuously performed and recorded every 10 s.

### Generation and cultivation of Arabidopsis plants

Transformation of *Arabidopsis thaliana* Col-0 plants for stable expression of NAPstar constructs was performed by floral dip [72]. Transformants were selected based on hygromycin resistance and sensor protein fluorescence. We isolated between 5–12 independent lines for each of the constructs. At least 3 lines with bright fluorescence were taken forward to homozygosity. The sensor fluorescence in the leaf epidermis was validated for all selected lines to show the characteristic cyto-nuclear pattern, a high signal-to-noise ratio when compared to the wild-type control, and clear separation of the biosensor fluorescence from chlorophyll autofluorescence. Other plant biosensor lines were used as reported previously; Peredox [23], Grx1-roGFP2 [56], roGFP1-Orp1 [73] and NERNST [18]. Plants were cultivated on soil made up from 50% VMV800 Vermehrungserde and 50% EDE800 Einheitserde (Balster Einheitserden, Frödenberg, Germany) at long-day conditions (16 h light, 8 h dark; light period: 100 to 150 µmol photons m^-2^ s^-1^ by OSRAM HO 54W/840 LUMILUX Cool White tubes; 22°C during light and 18°C during darkness; 65% humidity). After sowing, seeds were stratified at 4°C for 2–3 days before pots were moved into the cultivation chambers. Pots were watered from the bottom every 3–4 days.

### *In planta* biosensor measurements

The multiparametric real-time measurements were conducted with a CLARIOstar multiwell plate-reader equipped with an atmospheric control unit (BMG Labtech, Ortenberg, Germany). Leaf disc samples were cut from approximately five-week-old Arabidopsis rosettes with a leaf disc cork borer as described previously [23]. The leaf discs were transferred to wells of a NUNC 96-well plate (VWR International GMBH, Darmstadt, Germany) each filled with 200 µL 10 mM MES, 10 mM MgCl_2_, 10 mM CaCl_2_, 5 mM KCl, pH 5.8. Experiments were executed at 25°C in the early afternoon. The dark-light transition experiments were carried out as described previously [23]. The plate was removed from the plate reader for the duration of the illumination and placed back immediately after. An intensity of 600 µE was used for white light illumination with ELRO LED 1 x7 W IP44 (ELRO Europe, Amsterdam, Netherlands).

The hypoxia assays were performed as described previously [36, 37, 67, 74] with several adjustments. The sensor fluorescence response was measured over a time course and an oxygen ramping script mode together with the ACU was used to apply an oxygen gradient by flushing the measurement chamber with nitrogen gas (99.998 % v/v, Westfalen AG, Münster, Germany). The script mode was used to set a standardized measurement regime of 2.5 h at normoxic conditions, followed by 0.5 h oxygen depletion down to a defined hypoxic O_2_ concentration and a hypoxic phase of 6 h, followed by reoxygenation with ambient air within 0.5 h. The fluorescence was then recorded for another 7.5 h. The fluorescence readout of the sensors was recorded using the following excitation and emission bands: Peredox, NAPstar4.3 and NAPstarC: TS: Ex = 400 ±5 nm, Em = 520 ±5 nm; mCherry: Ex = 540 ±10 nm, Em = 615 ±9 nm; Grx1-roGFP2/roGFP2-Orp1, Ex = 400 ±5 nm and 482 ±8 nm, Em = 520 ±5 nm). Detector gain was set to allow optimal detection range while avoiding detector overflow and was kept constant for all measurements within an experiment. The emission was recorded using the top optic detector and the orbital averaging mode (diameter: 3 mm and 30-35 light flashes).

Background and autofluorescence were corrected by subtracting an emission average from wild type Col-0 tissue measured in parallel using the same treatments but without sensor expression. Biosensor fluorescence ratios were calculated as follows: Peredox and NAPstar family (TS/mCherry), and roGFP2 family (400/482 nm). The data were then log10-transformed. The data were normalized to zero by subtracting an average of the last five intensity values before oxygen depletion or before treatment from each time point value.

### Fluorescence lifetime imaging microscopy (FLIM)

For *in vitro* lifetime measurements, 0.025 µg µL^-1^ NAPstar4.3 diluted in binding buffer, containing 150 µM NADP^+^ was added to imaging 8-wells (80806; ibidi, Gräfelfing, Germany). Increasing amounts of NADPH (1-250 μM) were subsequently added to the wells and the solutions imaged on a LSM880 confocal laser scanning microscope (Carl Zeiss AG, Oberkochen, Germany) with a 63x immersion objective (LD LCI Plan-Apochromat 63x/1.2 Imm Corr DIC M27) using a 440 nm pulsed excitation laser (LDH-D-C-440, 40 MHz repetition rate) managed by a PicoQuant FLIM module (Sepia PDL828-S, PMA Hybrid 40, MultiHarp 150 4N, Time Harp 260 PICO Dual) (PicoQuant, Berlin, Germany). Fluorescence lifetime images of 512 x 512 pixels (135.2 μm x 135.2 μm) and 2 cycles or 5 x 10^5^ photon counts at a pixel dwell time of 16.38 μs passing through a 550 ±49 nm bandpass filter (Semrock, F37-551, IDEX Health & Science, LLC, Rochester, NY, United States) were recorded using the handshake plugin between Zen Black imaging software (Carl Zeiss AG, Oberkochen, Germany) and SymPhoTime 64 software (PicoQuant, Berlin, Germany). Ten FLIM images were recorded per treatment and measurements were repeated over three experimental days. Each set of images was analyzed using the ‘Grouped FLIM’ analysis in SymPhoTime 64 with a bi-exponential fitting model. The obtained amplitude weighted averaged lifetime, τ_Av Amp_, values were averaged for the set and used to plot the K_d_ curves.

### Cell culture and transient transfection

HeLa cells (Leibnitz Institute DSMZ; ACC 57) at passage 4-25 were cultured in Dulbecco’s Modified Eagle Medium (DMEM) (Gibco, Ref: 41966029, Thermofisher Scientific,Waltham, Massachusetts, United States) supplemented with 10% fetal bovine serum (FBS) (Gibco, Ref: 10270106, Thermofisher Scientific,Waltham, Massachusetts, United States) in humidified air at 37°C and 5% CO_2_. Cells were subcultured twice a week at a ratio of 1:5 to 1:8.Transient transfection of HeLa cells (1–2 x 10^5^ cells/6-well) with pcDNA3.1(+) NAPstar or pSC2 HyPer7 (Addgene; plasmid #136466, Watertown, MA, United States) plasmids were performed with the transfection reagent FuGENE® HD (Promega, Ref: E2311, Promega GmbH, Walldorf, Germany) at a 2:1 FuGENE® HD:DNA ratio according to the manufacturer protocol. After 48 h incubation at regular culture conditions the cells were used for fluorescence microscopy measurements.

### Fluorescence microscopy

#### Epifluorescence microscopy

NAPstar or HyPer7 expressing HeLa cells were analyzed with a Zeiss Axio Observer 7 inverted epifluorescence microscope and the ZEN3.2 blue edition software (Carl Zeiss AG, Oberkochen, Germany). NAPstar expression levels in HeLa and yeast cells were imaged using following excitation and emission settings: Ex: 405 ±20 nm, Em: 525 ±50 nm and Ex: 572 ±25 nm, Em: 629 ±62 nm. Yeast cell were imaged using a 100x objective, HeLa cells were imaged using a 63x objective. Cells were imaged in glucose-free HBSS. Images were analyzed using ImageJ software.

For perfusion assays with HeLa cells, NAPstar3b was monitored using Ex and Em settings as above, HyPer7 was monitored using- Ex: 405 ±20 nm, Em: 525 ±50 nm and Ex: 470 ±40 nm, Em: 525 ±50 nm; Cells were preincubated in glucose-free or 10 mM glucose containing Hank’s Balanced Salt Solution (HBSS) (1.3 mM CaCl_2_, 0.8 mM MgSO_4_, 5.4 mM KCl, 0.44 mM KH_2_PO_4_, 4.2 mM NaHCO_3_, 137 mM NaCl, 0.34 mM Na_2_HPO_4_ pH 7.4) for 1 h at 37°C and 5% (v/v) CO_2_. The measurement was performed in HBSS containing 0 mM or 10 mM glucose supplemented with H_2_O_2_ (2.5-20 µM) or menadione (1-10 µM) and fresh buffer solution were continuously provided by a peristaltic pump adjusted to a flow speed of 1 mL min^-1^.

#### Confocal microscopy

A Zeiss LSM980 confocal microscope operated using the ZEN3.6 blue edition software (Carl Zeiss AG, Oberkochen, Germany) was used for plant cell imaging. A 40x C-Apochromat lens with 1.2 numeric aperture and water immersion was used. The pinhole was set in range of 1-2 airy units. Leave disks used for microscopy were at least incubated in the dark for 2 h. The TS fluorophore and chlorophyll autofluorescence were excited at 405 nm and emitting light was measured between 497 nm and 542 nm (TS) and in between 654 nm and 701 nm (Chlorophyll). The mCherry fluorophore was excited at 561 nm and the emission was collected between 588 nm and 632 nm.

### Generation of CRISPR KO strains

To generate a knockout of glutathione reductase (GSR) in HEK293 cells, a guide RNA sequence targeting exon 1 was cloned into the pSpCas9(BB)-2A-GFP (PX458) vector, gifted from Feng Zhang (Addgene plasmid #48138, Watertown, MA, United States) [75]. HEK flp-InTm T-RExTM-293 cells (Invitrogen, Darmstadt, Germany) were transfected with this construct using a standard polyethylenimine-based approach. 24 h after transfection, cells were sorted by FACS, based on GFP fluorescence. Cells were then seeded on a single cell basis into 96-well plates and screened for loss of protein using western blot analysis (Proteintech, Cat. Number: 18257-1-AP, GSR Rabbit polyclonal antibody).

### Cytation measurements

High-throughput imaging measurements of HEK293 cells expressing NAPStar3b from a pcDNA3.1(+) plasmid and data analysis were performed as previously described [76]. Experiments were performed using wildtype and GSR KO HEK293 cells. In each well of a poly-L-lysine coated 96-well plate (μclear, GreinerBio, Kremsmünster, Austria), approximately 4,000 cells were seeded in 100 µl Dulbecco’s Modified Eagle Medium (DMEM) containing 10% fetal calf serum (FCS) and 1% penicillin and streptomycin (P/S). 24 h after seeding, cells were transfected with the plasmid containing the indicated construct using a standard polyethylenimine-based method.

After 48 h, the measurement was started by switching from DMEM medium to preheated glucose starvation minimal medium (140 mM NaCl, 5 mM KCl, 1 mM of MgCl_2_, 2 mM of CaCl_2_, 20 mM Hepes, pH 7.4 as adjusted with NaOH) with 10% FCS. Subsequently, the plate was incubated in the Cytation3 (BioTek, Winooski, VT, United States) at 37°C with 5% (v/v) CO_2_ for 30 min. The sensor fluorescence was measured using the 10x air objective. A 405 nm and 590 nm BioTek filter cube was used for readouts of either 400 ±40 nm or 586 ±15 nm respectively. After the incubation step, the steady state was monitored for 30 min. For inhibition of thioredoxin reductase or glutathione reductase activity, cells were subsequently treated with 1 µM auranofin (Sigma Aldrich, Darmstadt, Germany) or 50 µM 1,3-bis(2-chloroethyl)-1-nitrosourea (BCNU) (Sigma Aldrich, Darmstadt, Germany) respectively for 1 h. Following the treated with 50 and 100 µM H_2_O_2_ or 100 and 500 µM diamide, sensor responses were monitored for 1 h. Data analysis was performed using the RRA custom Matlab analysis suite [77] and 490 nm over 585 nm ratios were calculated.

