## Supplementary Figures for "The NAPstar family of NADP redox state sensors highlights glutathione as the primary mediator of anti-oxidative electron flux"

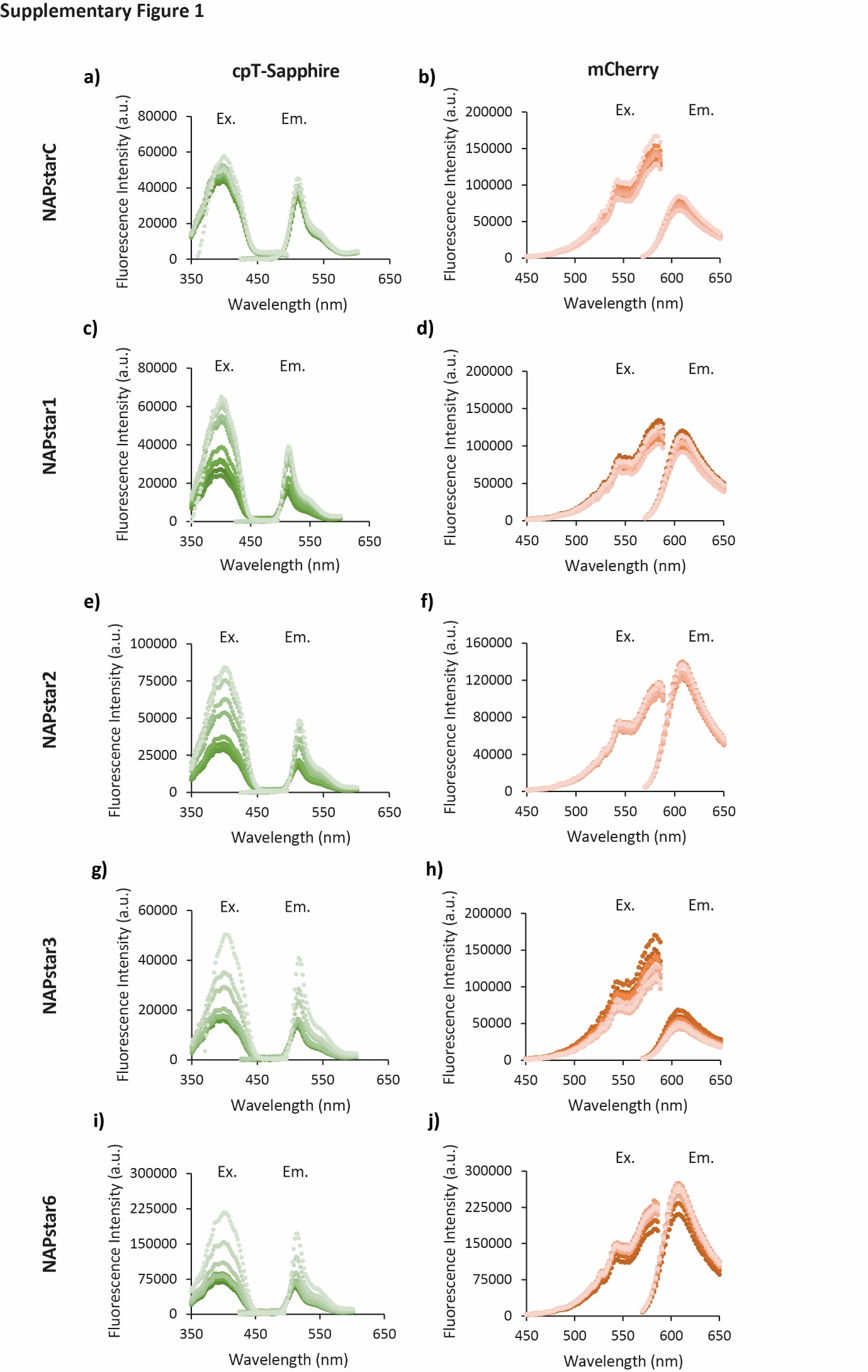

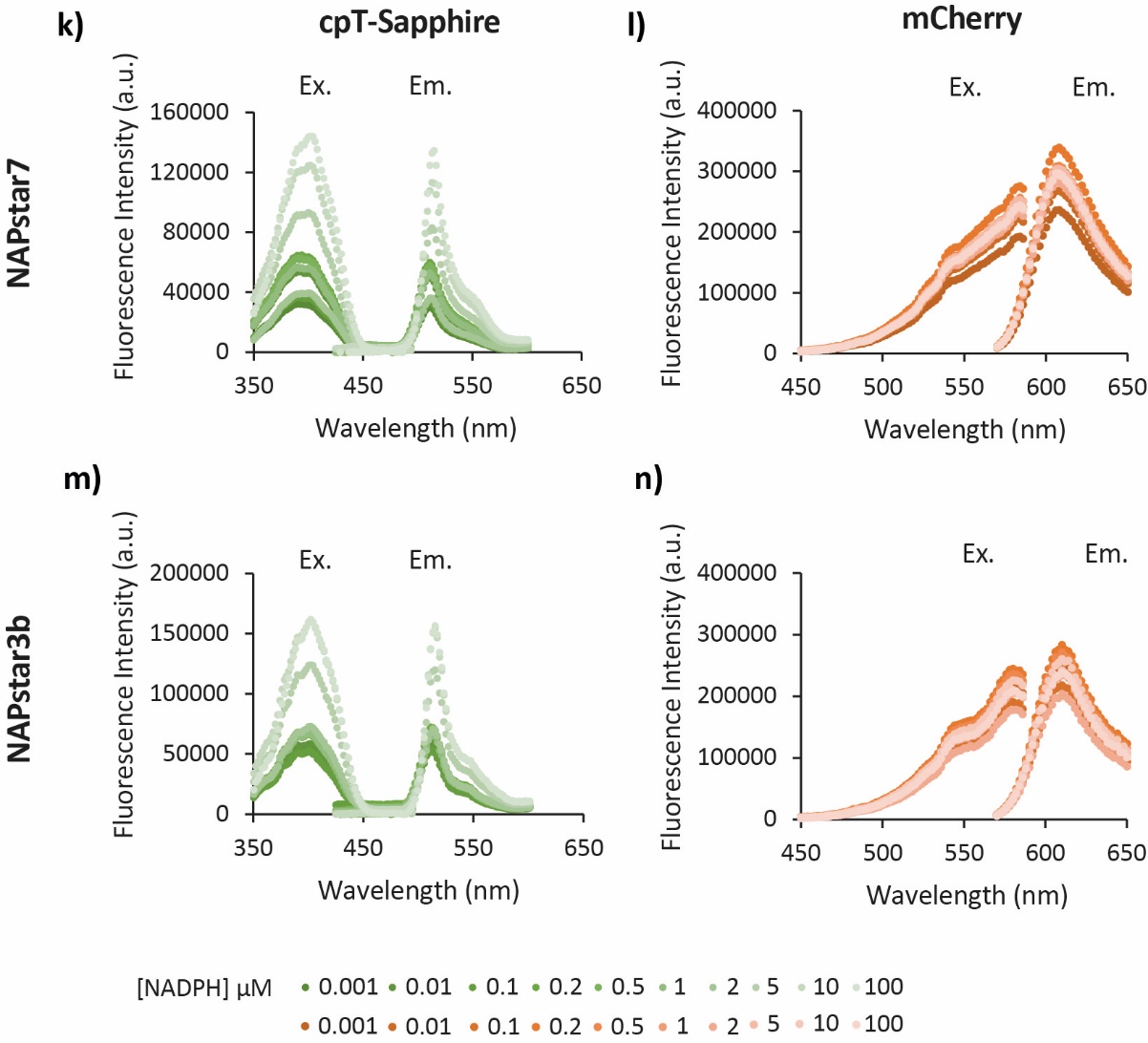

**Supplementary Figure 1. Fluorescence spectra of NAPstars.**

Fluorescence excitation (Ex.) and emission (Em.) intensity spectra of (**a,c,e,g,i,k,m**) the cpT-Sapphire and the mCherry (**b,d,f,h,j,l,n**) fluorophores of the different NAPstar variants, incubated in the presence of 150 µM NADP^+^ and NADPH at the indicated concentrations ranging from 0–100 µM. Sensor protein concentrations were adjusted to 240 nM (n=3 technical replicates). Data are presented as mean ± s.d.

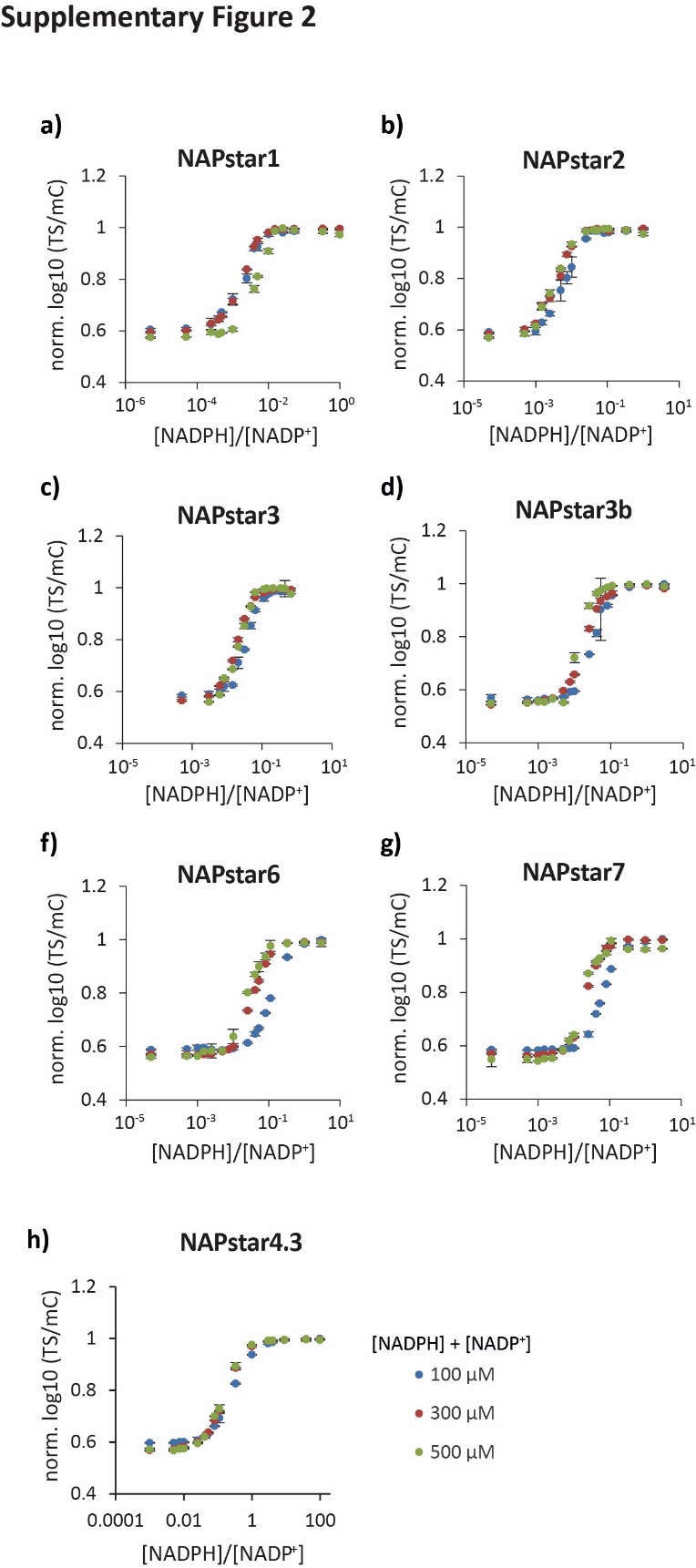

**Supplementary Figure 2. NAPstars report the NADP redox state.**

**a–h,** NAPstar variants were incubated in the presence of total NADP concentrations ([NADPH] + [NADP^+^]) of 100 µM, 300 µM and 500 µM. Different NADP redox states were set by adjustment of the relative NADPH and NADP^+^ concentrations. Sensor protein concentrations were adjusted to 240 nM for all measurements (n=3 technical replicates). Data are presented as mean ± s.d.

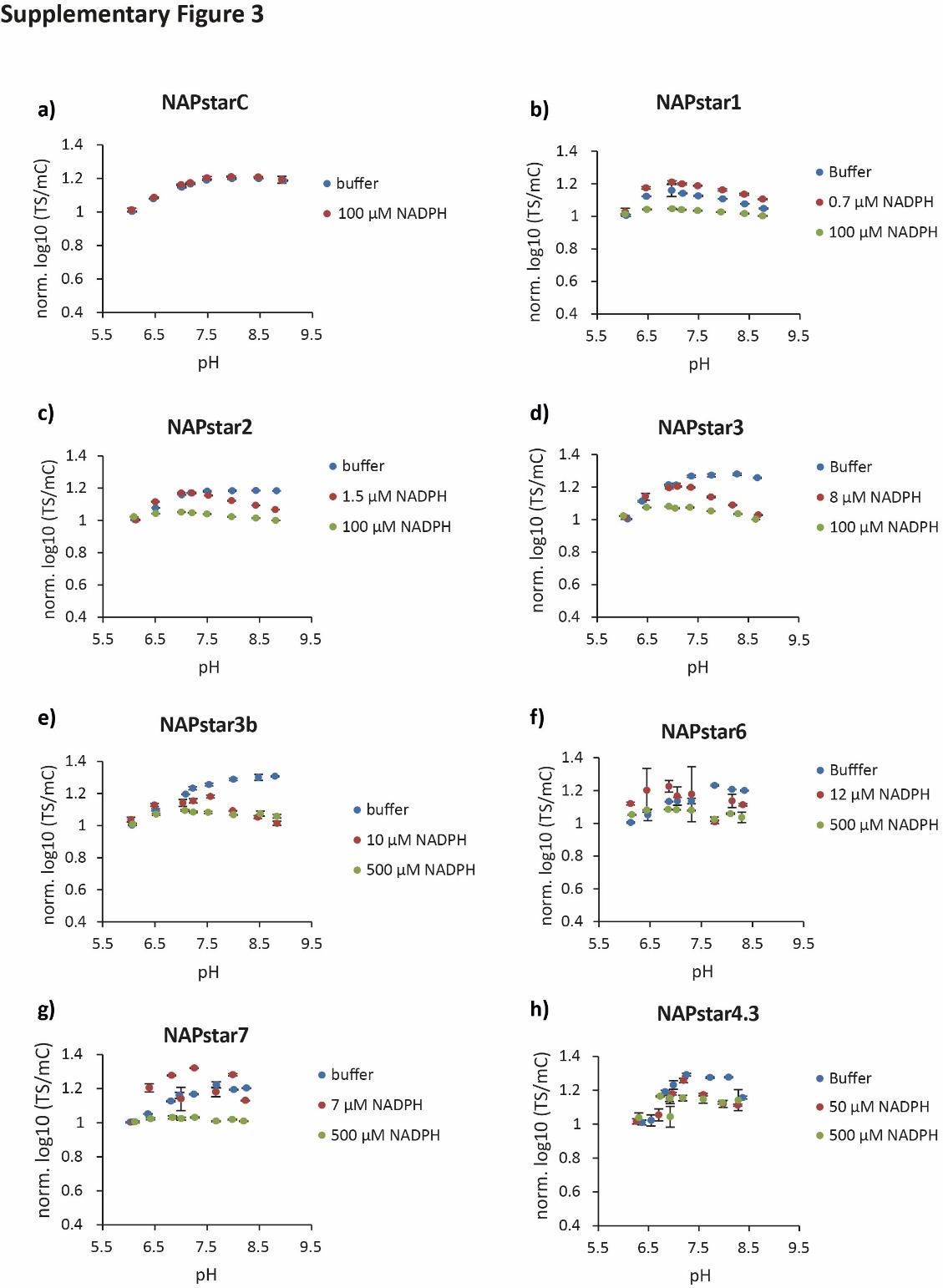

**Supplementary Figure 3. NAPstar measurements are resistant to pH changes.**

**a–h**, Individual NAPstar family proteins were incubated in buffers with different pH values spanning the physiologically relevant pH range (see *Materials & Methods* section). All probes were incubated in the presence of 150 µM NADP^+^ and NADPH at the indicated concentrations to establish different NADPH or NADP^+^ occupancies. NADPH concentrations were used to establish saturation by NADP^+^ binding only (buffer; i.e. no NADPH; blue), approximately half occupancy by NADP^+^ and NADPH each as informed by the the specific *K*_r_ of the respective NAPstar variant (red), and saturation by NADPH binding only (100 or 500 µM NADPH; green). An exception is NAPstarC (**a**) due to lack of substrate binding. Sensor protein concentrations were adjusted to 240 nM (n=3 technical replicates for all measurements). Data are presented as mean ± s.d.

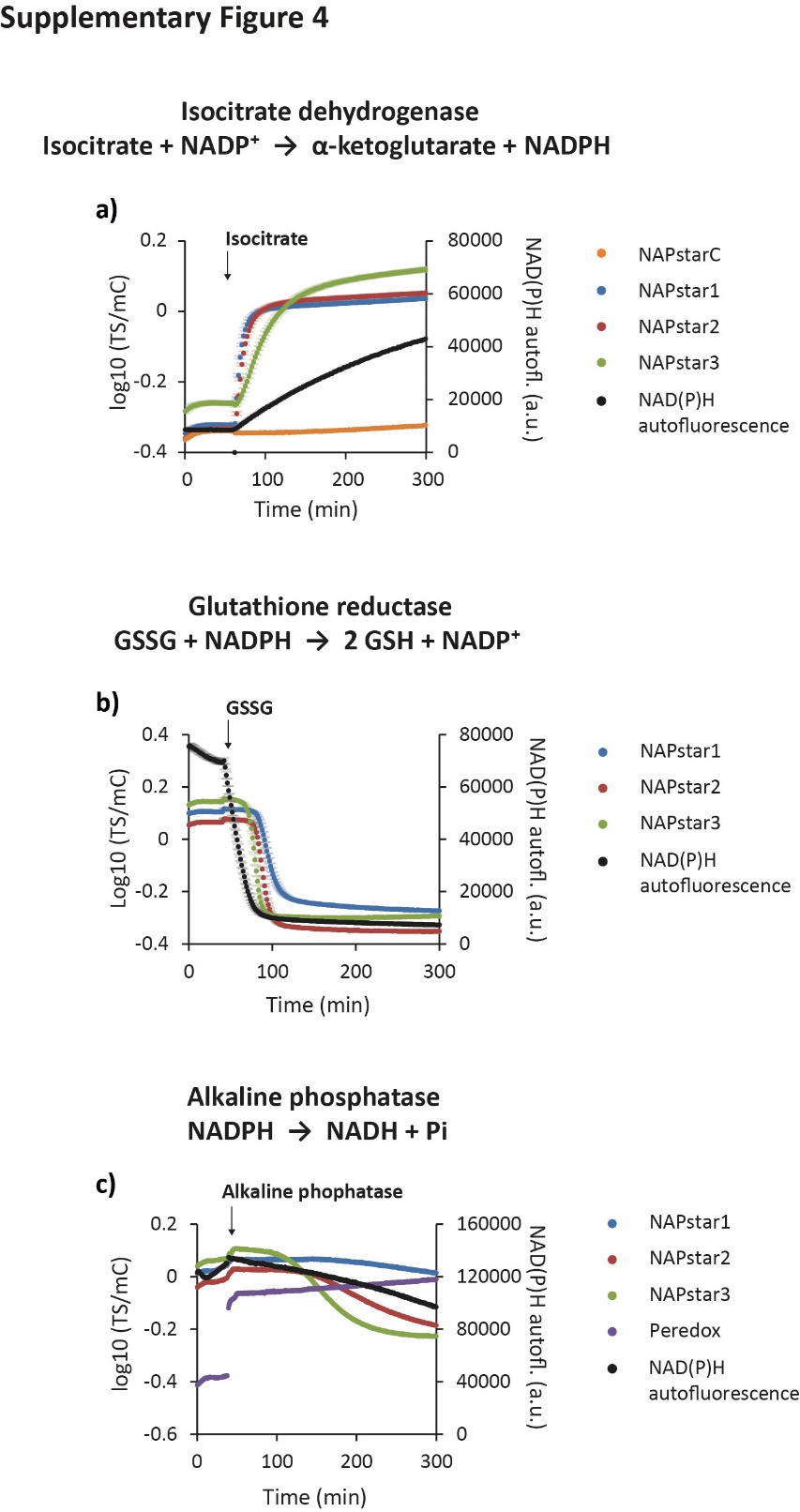

**Supplementary Figure 4. NAPstar-based online monitoring of NADP redox dynamics in vitro.**

**a,** NADP^+^ (initial concentration 500 µM) to NADPH conversion was initiated by the addition of 1 mM isocitrate in the presence of isocitrate dehydrogenase. **b,** NADPH (initial concentration 100 µM) to NADP^+^ conversion was initiated by the addition of 1 mM glutathione disulfide in the presence of glutathione reductase. **c,** NADPH (initial concentration 100 µM) to NADH conversion was initiated by the addition of alkaline phosphatase. Enzyme origin, quantities and assay buffer conditions are specified in the *Materials and Methods* section. Sensor protein concentrations were adjusted to 430 nM (n=3 technical replicates **a,b** or one replicate, **c**). Data are presented as mean ± s.d.

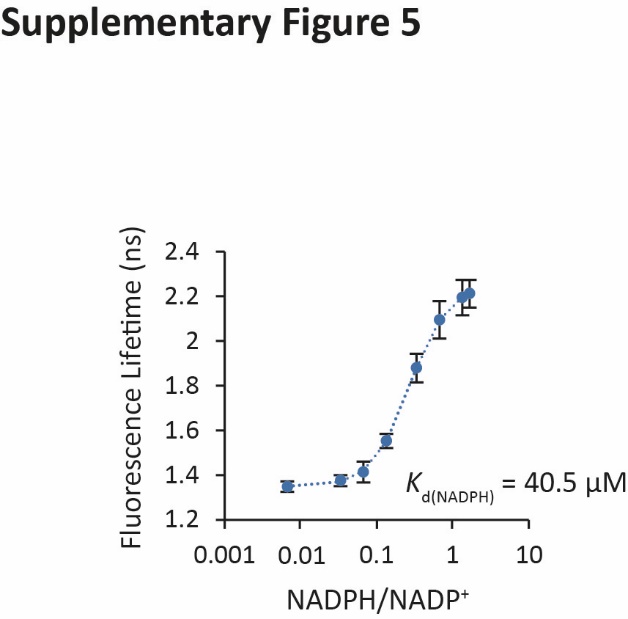

**Supplementary Figure 5. NAPstars are well suited to fluorescence-lifetime measurement.**

**a**, NAPstar4.3 was incubated in the presence of 150 µM NADP^+^ and different NADPH/NADP^+^ ratios were established by adjusting NADPH (1-250 μM). Fluorescence lifetime measurements of cpT-Sapphire signal were performed at 440 nm pulsed laser excitation and 550/49 nm bandpass filter emission as detailed in the *Materials & Methods* section. Sensor protein concentrations were adjusted to 240 nM. *K*_d(NADPH)_ was calculated at 40.5 µM. (n=3 independent experiments, each with 10 technical replicates). Data are presented as mean ± s.d.

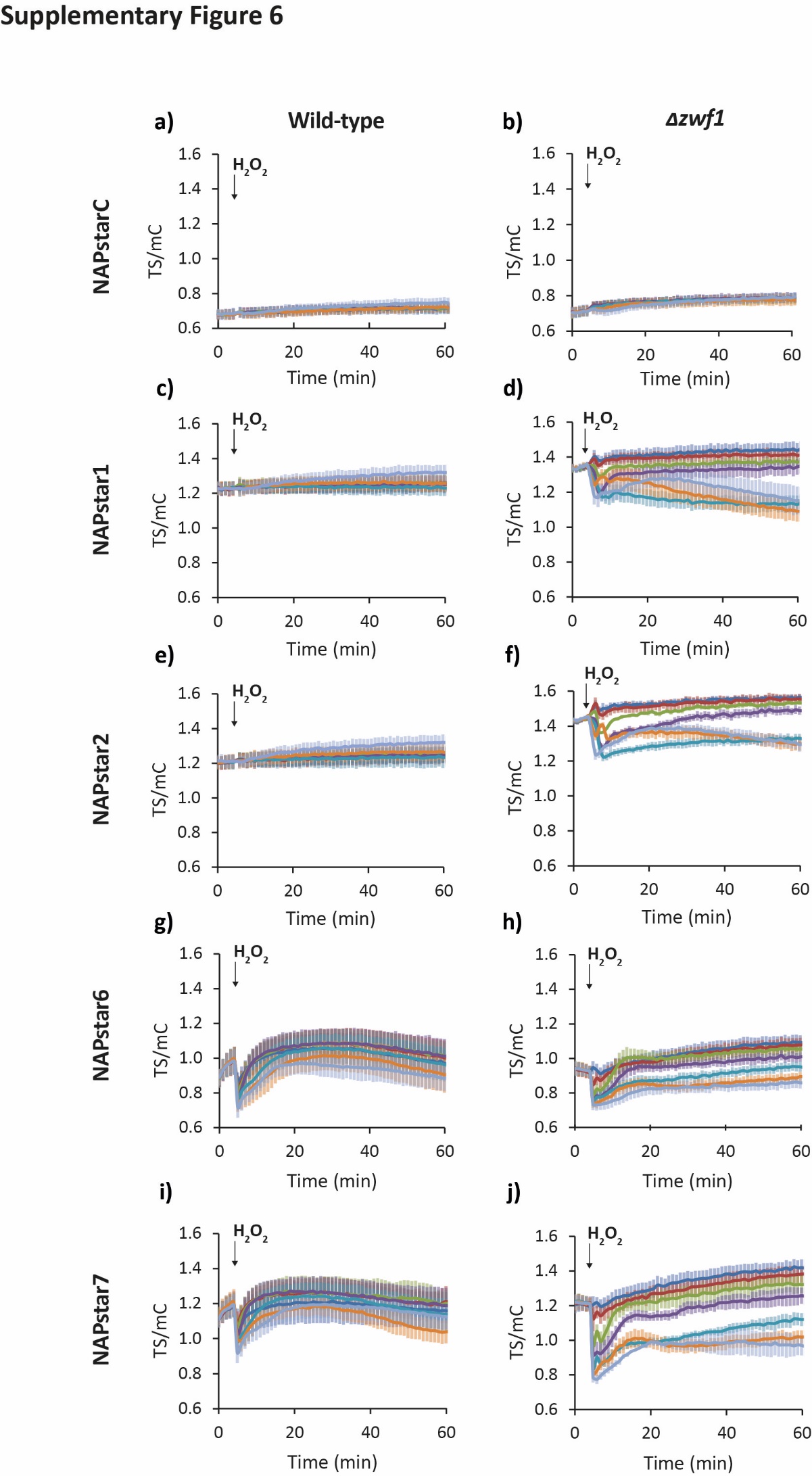

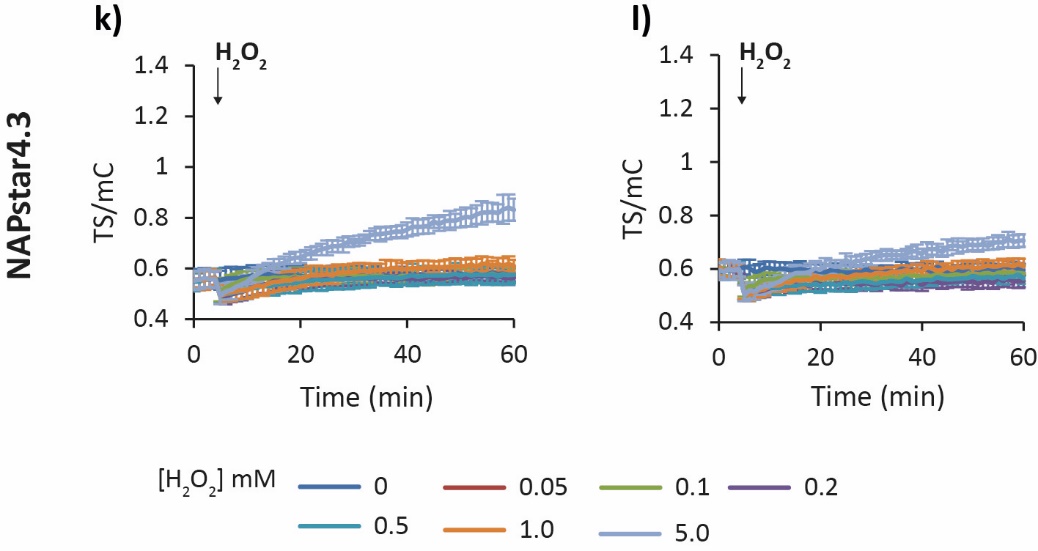

**Supplementary Figure 6. NAPstar responses in the yeast cytosol to external H_2_O_2_.**

Related to **Figure 2**. Response of wild-type (**a, c, e, g, i, k**) and *Δzwf1* (**b, d, f, h, j, l**) yeast cells expressing the indicated NAPstar variants, to treatment with H_2_O_2_ at the indicated concentrations (n=3 repeats with cells from independent cultures). Data are presented mean ± s.d.

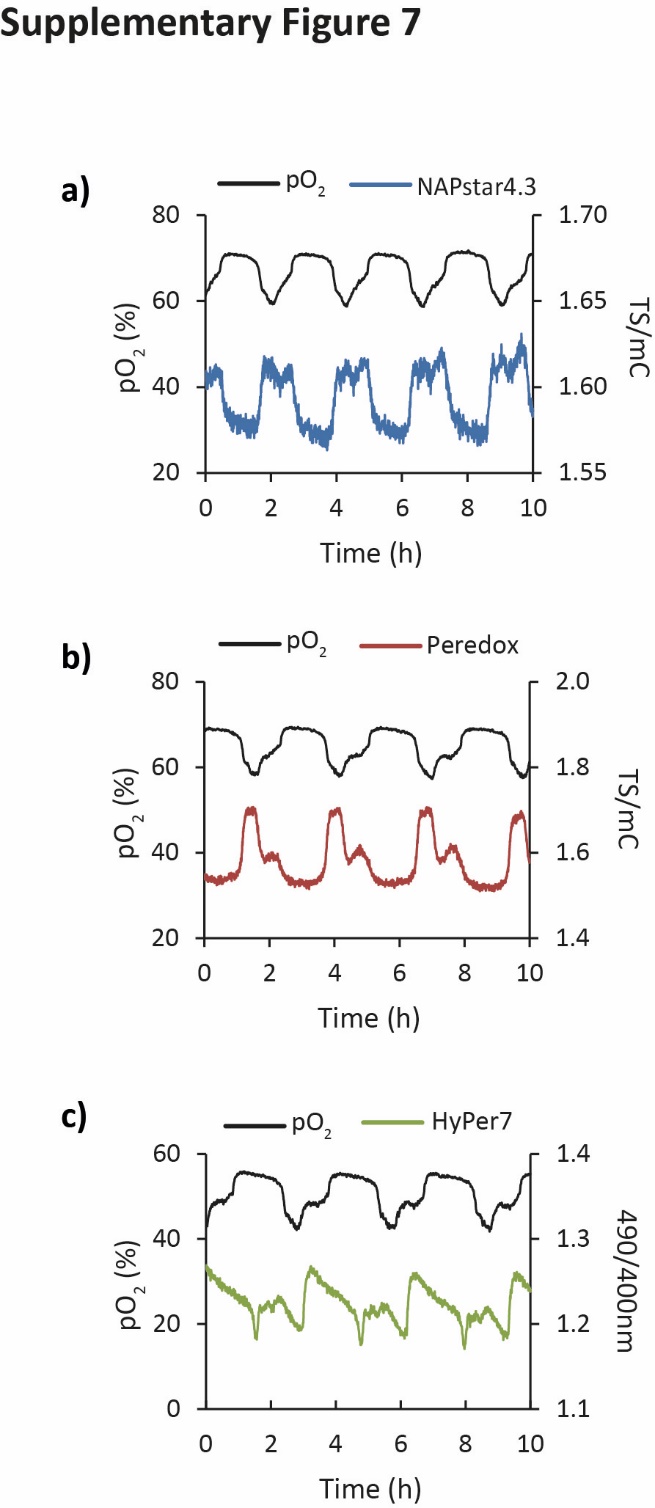

**Supplementary Figure 7. Oscillations in cytosolic NADP and NAD redox state accompany the yeast metabolic cycle (YMC).**

Related to **Figure 3**. **a–c,** Replicate traces supporting **Fig. 3c**, showing the changes in dissolved oxygen, NAPstar4.3 (NADP redox state), Peredox (NAD redox state) and Hyper7 (H_2_O_2_) during 3.5 complete cycles of the YMC.

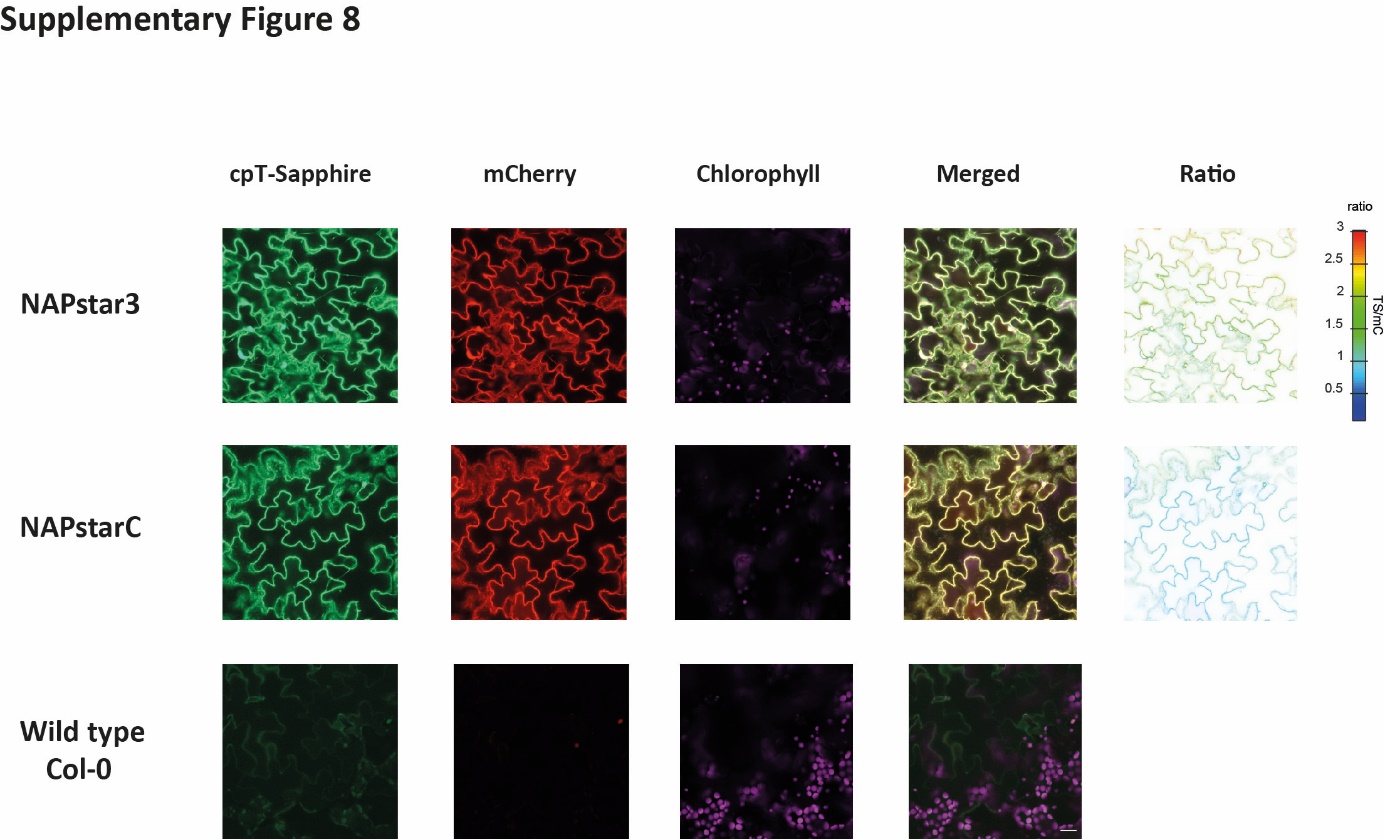

**Supplementary Figure 8. NAPstar expression in Arabidopsis leaf epidermis.**

Confocal microscopy images of NAPstar3, NAPstarC and the wild-type (Col-0) in the cytosol of *Arabidopsis thaliana* plants. Images show the abaxial leaf epidermal layer. Identical microscope settings were used for all lines (see *Material & Methods*). Corresponding ratio images that display TS/mC are shown on the right as calculated using the RRA custom software package (see *Material & Methods*). Scale bar = 20 µm.

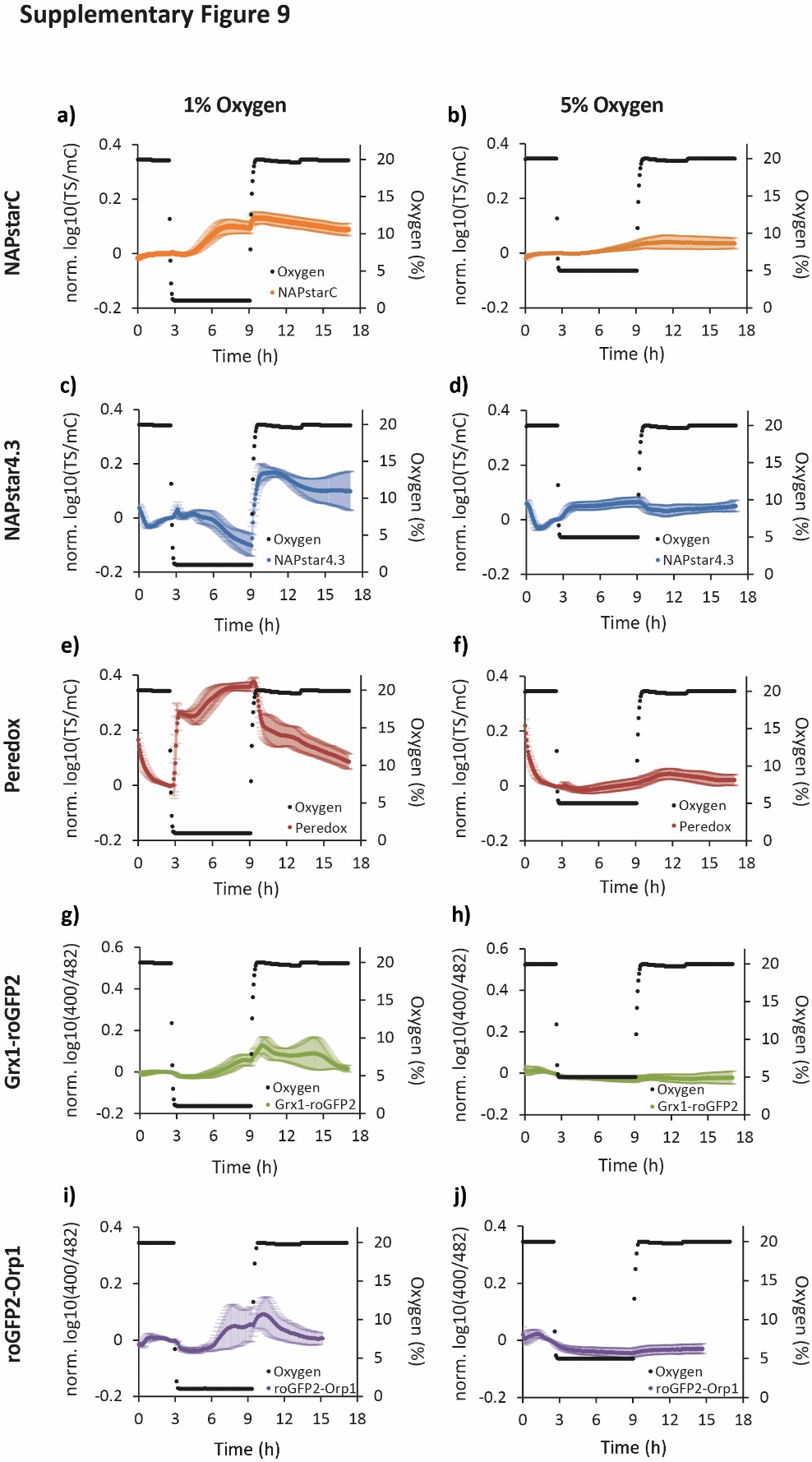

**Supplementary Figure 9. Cytosolic redox dynamics accompany hypoxia-reoxygenation in plants.**

Related to **Figure 4**. **a,b,** NAPstarC (control), **c,d,** NAPstar4.3 (NADP redox state), **e,f,** Peredox (NAD redox state), **g,h,** Grx1-roGFP2 (glutathione redox potential; *E*_GSH_), and **i,j,** roGFP2-Orp1 (H_2_O_2_) dynamics in response to the indicated period of hypoxia with 1% and 5% oxygen respectively (n=6–8 leaf discs from individual plants). In all panels, data are presented as mean ± s.d.

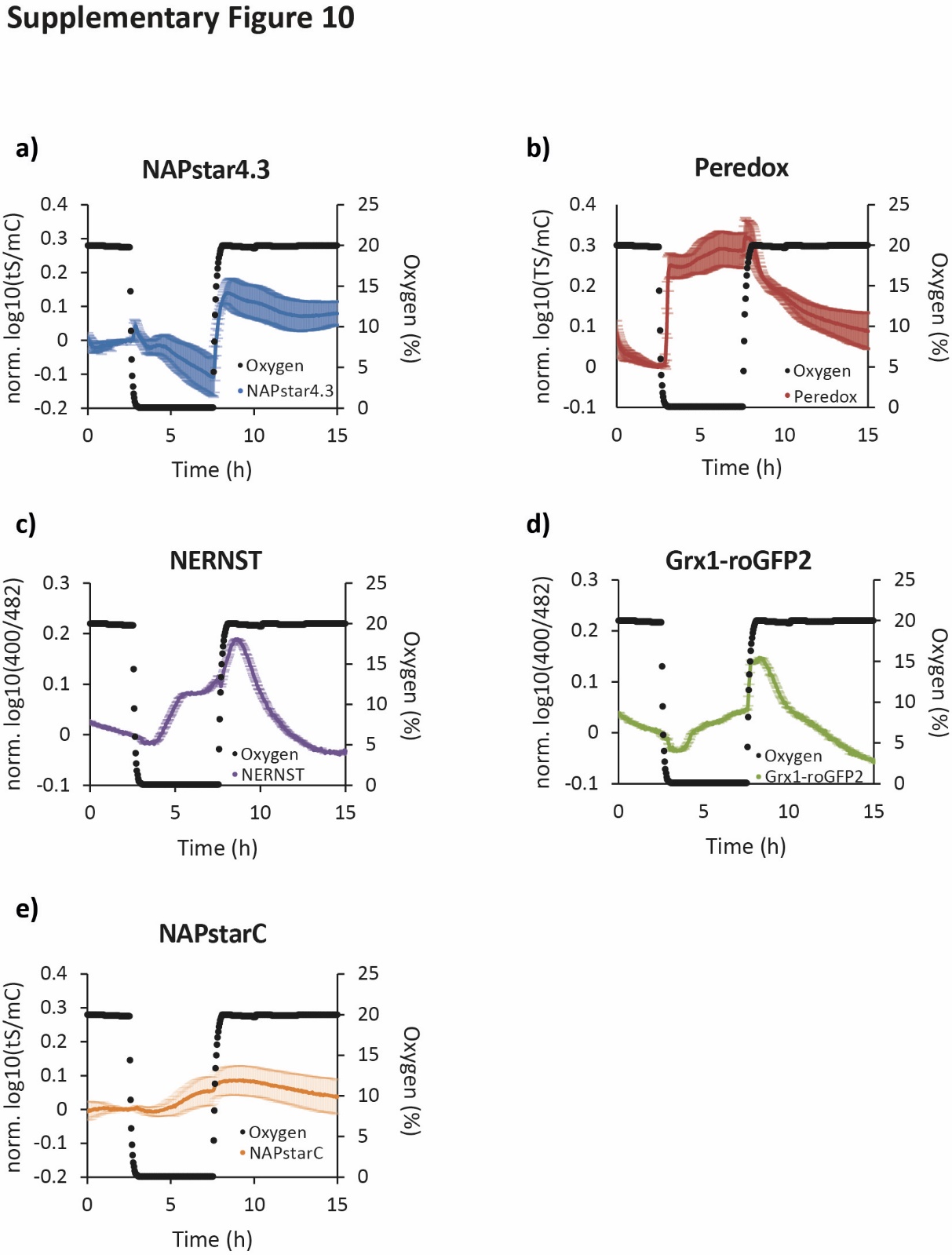

**Supplementary Figure 10. NERNST resembles the dynamics of Grx1-roGFP2 but not NAPstar4.3 during hypoxia-reoxygenation in plants.**

Related to **Figure 4**. **a–e,** Response of NAPstar4.3, Peredox, NERNST; Grx1-roGFP2 and NAPstarC probes to hypoxia with 0.1% oxygen (n=4–6 leave discs from individual plants). In all panels, data are presented as mean ± s.d.

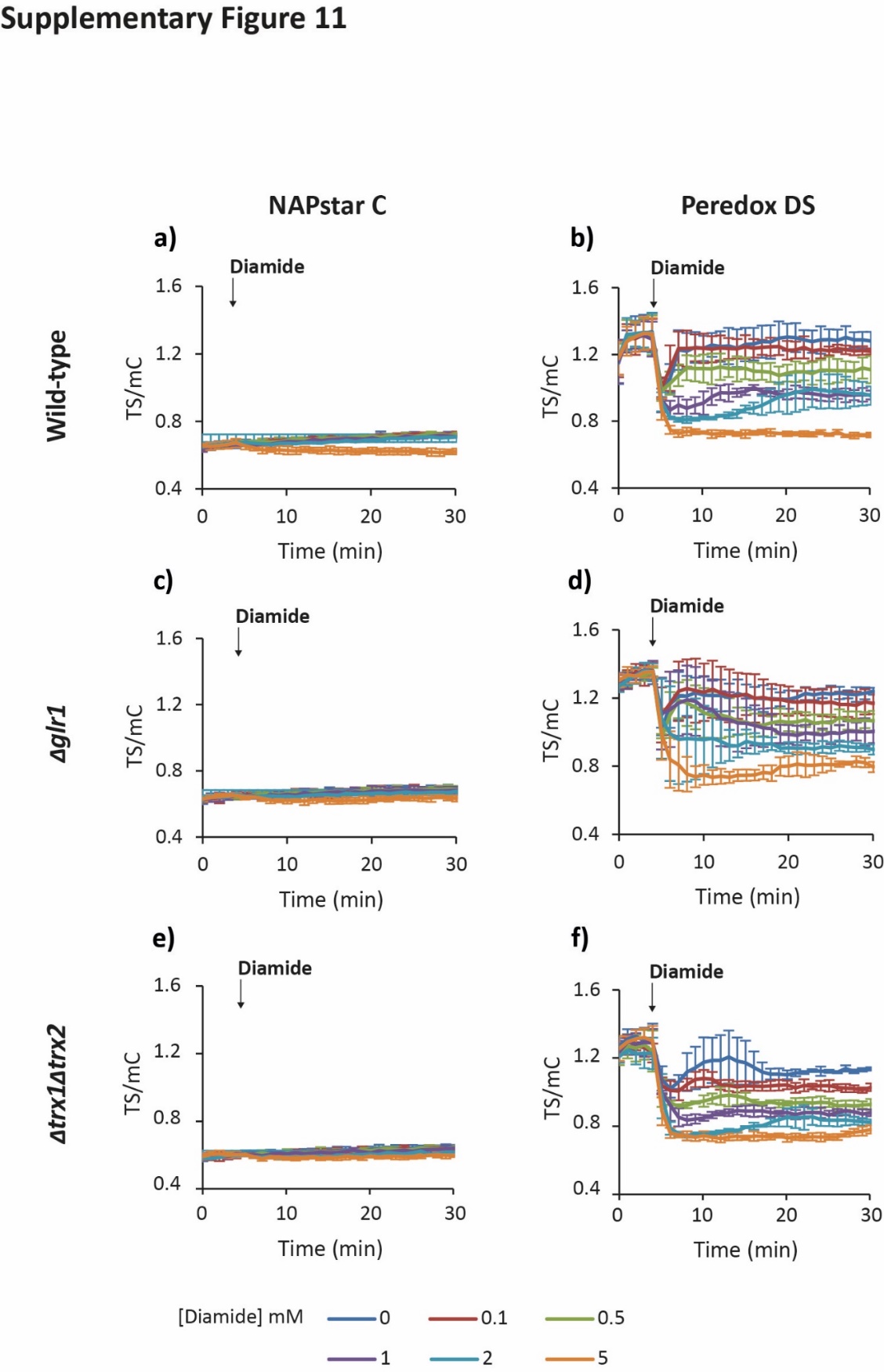

**Supplementary Figure 11. Glutathione reductase deletion does not change NAPstarC and PeredoxDS responses to diamide in yeast.**

Related to **Figure. 5**. Response of cytosolic NAPstarC or PeredoxDS probes to exogenous diamide at the indicated concentrations in wild-type (**a, b**), Δ*glr1* (**c, d**), and Δ*trx1*Δ*trx2* (**e, f**) cells (n=3 measurements made with cells derived from independent cultures). In all panels, data are presented as mean ± s.d.

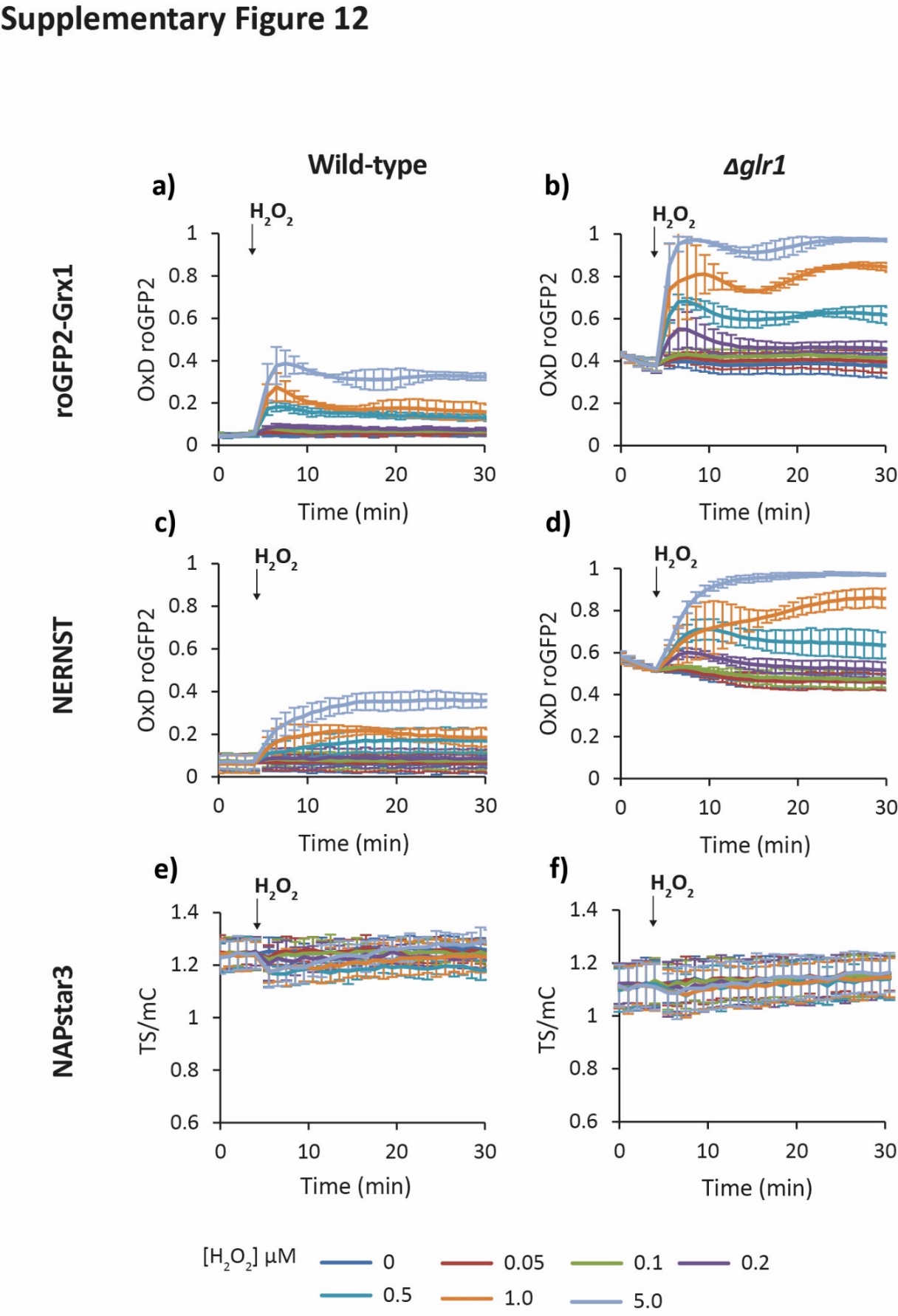

**Supplementary Figure 12. NERNST resembles the dynamics of Grx1-roGFP2 in yeast when the NADP and the glutathione redox systems are uncoupled by the absence of glutathione reductase.**

Related to **Figure 5**. Response of cytosolic roGFP2-Grx1, NERNST and NAPstar3 probes to exogenous H_2_O_2_ at the indicated concentrations in wild-type (**a, c, e**) and Δ*glr1* cells (**b, d, f**) (n=3 measurements made with cells derived from independent cultures). In all panels, data are presented as mean ± s.d.

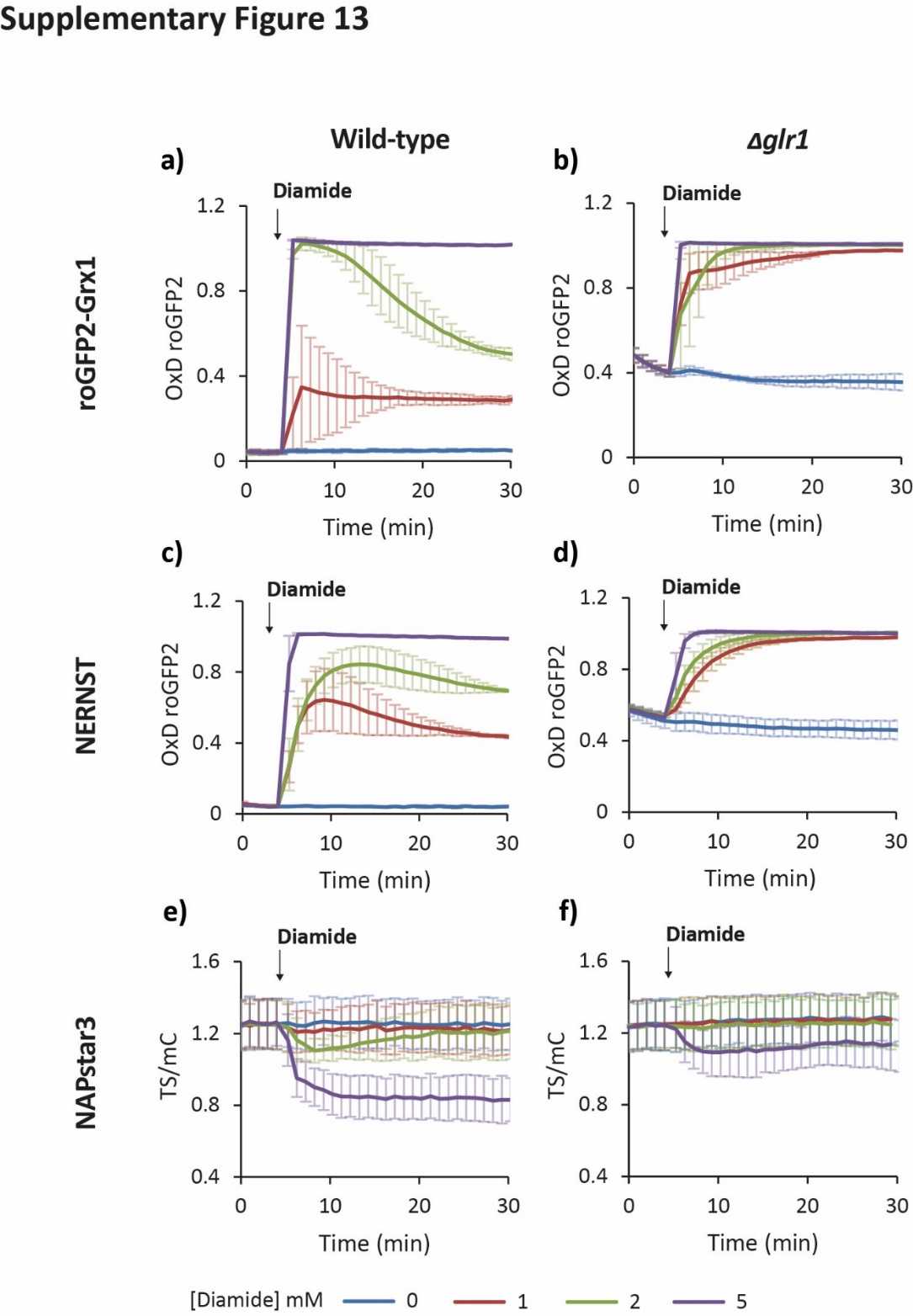

**Supplementary Figure 13. NERNST resembles the dynamics of Grx1-roGFP2 in yeast when the NADP and the glutathione redox systems are uncoupled by an absence of glutathione reductase.**

Related to **Figure 5**. Response of cytosolic roGFP2-Grx1, NERNST and NAPstar3 probes to exogenous diamide at the indicated concentrations in wild-type (**a, c, e**) and Δ*glr1* cells (**b, d, f**) (n=3 measurements made with cells derived from independent cultures). In all panels, data are presented as mean ± s.d.

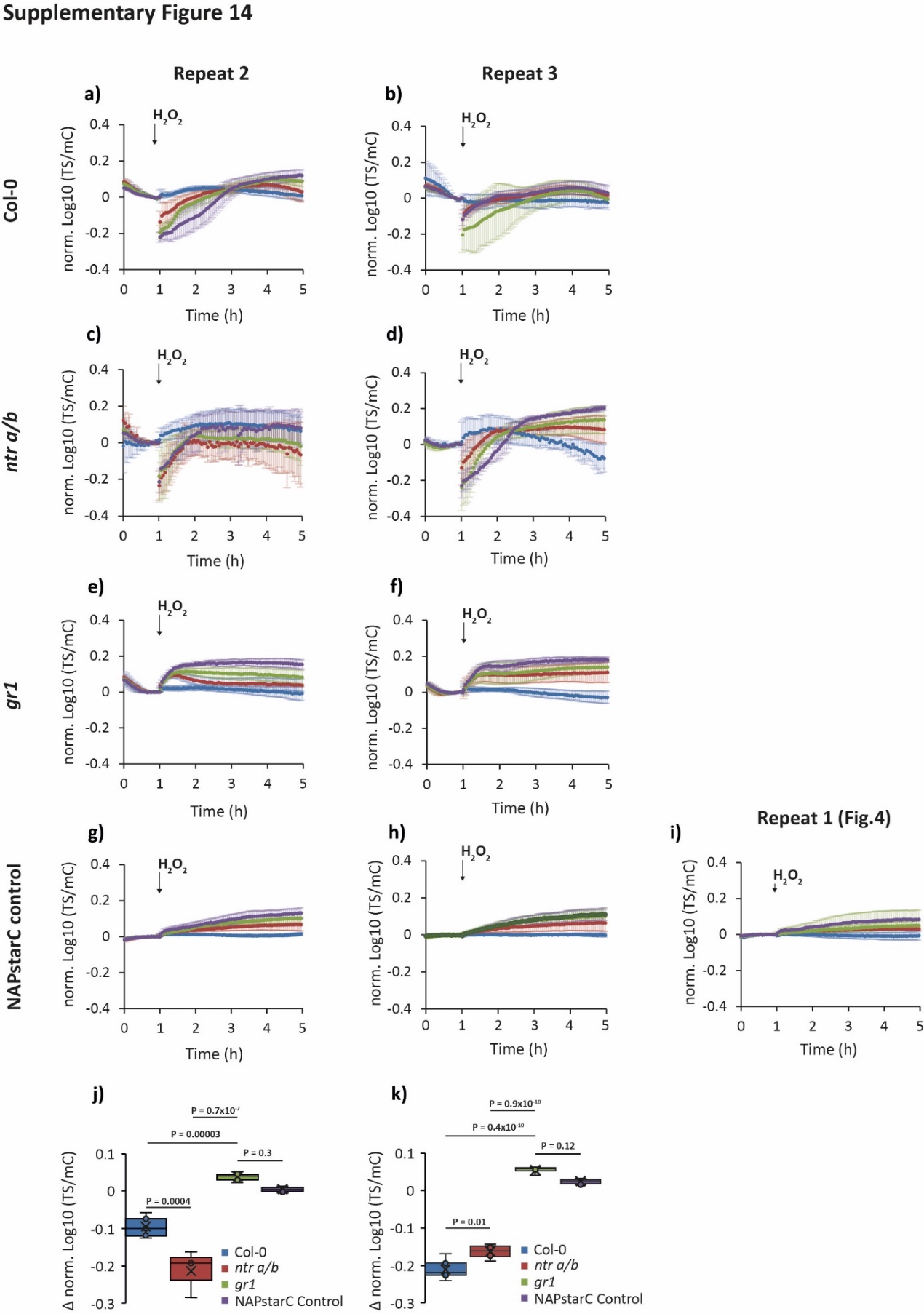

**Supplementary Figure 14. Glutathione reductase deletion protects against H_2_O_2_-induced NADP oxidation in plants.**

Related to **Figure. 6**. Two further experimental repeats, with the corresponding NAPstarC control responses, of the data in **Fig. 6b–e**.

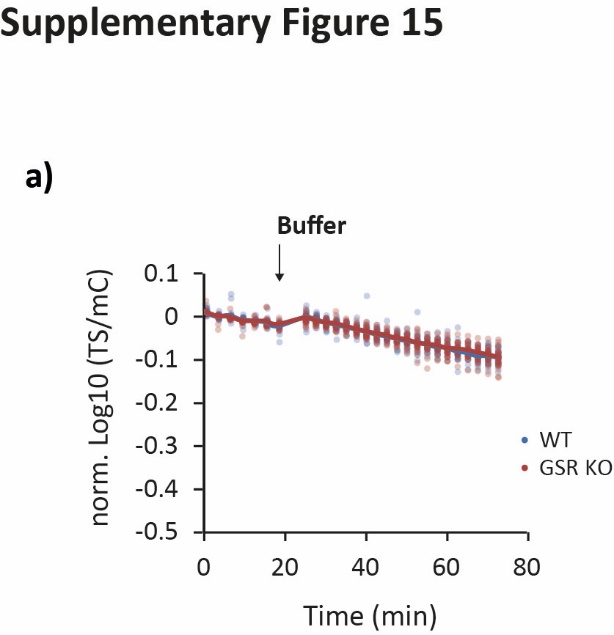

**Supplementary Figure 15. Glutathione reductase deletion protects against H_2_O_2_-induced NADP oxidation in HEK293 cells.**

Related to **Figure 7.** Response of NAPstar3b expressed in the cytosol of wild-type and glutathione reductase deleted (GSR KO) HEK293 cells to the buffer control (n=26 individual cells measured in two independent experimental repeats for both wild-type and GSR KO cells).
