## Supplementary Information for "The NAPstar family of NADP redox state sensors highlights glutathione as the primary mediator of anti-oxidative electron flux"

**Supplementary Table 1: Plasmids generated for cloning.**

| **Backbone** | **Sensor** | **Generated by** |
| --- | --- | --- |
| pDONR207 | NAPstarC | GeneScript Biotech, Rijswijk, Netherlands |
| pDONR207 | NAPstar1 | GeneScript Biotech, Rijswijk, Netherlands |
| pDONR207 | NAPstar2 | GeneScript Biotech, Rijswijk, Netherlands |
| pDONR207 | NAPstar3 | GeneScript Biotech, Rijswijk, Netherlands |
| pDONR207 | NAPstar4 | GeneScript Biotech, Rijswijk, Netherlands |
| pDONR207 | NAPstar3b | Site-directed mutagenesis |
| pDONR207 | NAPstar6 | GeneScript Biotech, Rijswijk, Netherlands |
| pDONR207 | NAPstar7 | GeneScript Biotech, Rijswijk, Netherlands |
| pDONR207 | NAPstar4.3 | *ApaI* and *HindIII* digest |

**Supplementary Table 2: Plasmids generated for plant expression.**

| **Backbone** | **Sensor** | **Generated by** |
| --- | --- | --- |
| pSS02 | NAPstarC | Gateway Cloning |
| pSS02 | NAPstar1 | Gateway Cloning |
| pSS02 | NAPstar2 | Gateway Cloning |
| pSS02 | NAPstar3 | Gateway Cloning |
| pSS02 | NAPstar3b | Gateway Cloning |
| pSS02 | NAPstar4 | Gateway Cloning |
| pSS02 | NAPstar6 | Gateway Cloning |
| pSS02 | NAPstar7 | Gateway Cloning |
| pSS02 | NAPstar4.3 | Gateway Cloning |

**Supplementary Table 3: Plasmids generated for bacterial expression.**

| **Backbone** | **Sensor** | **Generated by** |
| --- | --- | --- |
| pETG10a | NAPstarC | Gateway Cloning |
| pETG10a | NAPstar1 | Gateway Cloning |
| pETG10a | NAPstar2 | Gateway Cloning |
| pETG10a | NAPstar3 | Gateway Cloning |
| pETG10a | NAPstar3b | Gateway Cloning |
| pETG10a | NAPstar4 | Gateway Cloning |
| pETG10a | NAPstar6 | Gateway Cloning |
| pETG10a | NAPstar7 | Gateway Cloning |
| pETG10a | NAPstar4.3 | Gateway Cloning |

**Supplementary Table 4: Plasmids generated for mammalian expression.**

| **Backbone** | **Sensor** | **Generated by** |
| --- | --- | --- |
| pcDNA3.1(+) | HyPer7 | GeneScript Biotech, Rijswijk, Netherlands |
| pcDNA3.1(+) | NAPstar3b | GeneScript Biotech, Rijswijk, Netherlands |

**Supplementary Table 5: Plasmids generated for yeast expression.**

| **Backbone** | **Sensor** | **Generated by** |
| --- | --- | --- |
| p413TEF | NAPstarC | GeneScript Biotech, Rijswijk, Netherlands |
| p413TEF | NAPstar1 | GeneScript Biotech, Rijswijk, Netherlands |
| p413TEF | NAPstar2 | GeneScript Biotech, Rijswijk, Netherlands |
| p413TEF | NAPstar3 | GeneScript Biotech, Rijswijk, Netherlands |
| p413TEF | NAPstar4 | GeneScript Biotech, Rijswijk, Netherlands |
| p413TEF | NAPstar6 | GeneScript Biotech, Rijswijk, Netherlands |
| p413TEF | NAPstar7 | GeneScript Biotech, Rijswijk, Netherlands |
| p413TEF | NAPstar4.3 | Restriction digest/Ligation |
| p413TEF | Peredox | GeneScript Biotech, Rijswijk, Netherlands |
| p413TEF | PeredoxDS | Site-directed mutagenesis |
| p413TEF | NERNST | GeneScript Biotech, Rijswijk, Netherlands |
| p413TEF | HyPer7 | GeneScript Biotech, Rijswijk, Netherlands |
| p413TEF | roGFP2-Grx1 | *XbaI* and *XhoI* digest |

**Supplementary Table 6: Primers used in this study.**

| **Number** | **Primer** | **Sequence** | **Purpose** | **Company** |
| --- | --- | --- | --- | --- |
| Pr1 | NAPstar3b_for | TCGCGGTGGTTATATCGAACACGTCG | Generation of plant/ in vitro NAPstar3B | Merck KGaA, Darmstadt, Germany |
| Pr2 | NAPstar3b_rev | ACCGGCCGACCAACCTTC | Generation of plant/ in vitro NAPstar3B | Merck KGaA, Darmstadt, Germany |
| Pr3 | Peredox_D117S_fwd | GAATTGAGAGGTTTCTTTTCCGTTGATCCAGGCATGGTTGGTAGACC | Mutagenesis for generation of PeredoxDS | Eurofins Genomics GmbH, Ebersberg, Germany |
| Pr4 | Peredox_D117S_rev | GGTCTACCAACCATGCCTGGATCAACGGAAAAGAAACCTCTCAATTC | Mutagenesis for generation of PeredoxDS | Eurofins Genomics GmbH, Ebersberg, Germany |
| Pr5 | Peredox_D574S_fwd | CATTTGAATTAAGAGGTTTCTTTTCCGTTGATCCAGGCATGGTCGG | Mutagenesis for generation of PeredoxDS | Eurofins Genomics GmbH, Ebersberg, Germany |
| Pr6 | Peredox_D574S_rev | CCGACCATGCCTGGATCAACGGAAAAGAAACCTCTTAATTCAAATG | Mutagenesis for generation of PeredoxDS | Eurofins Genomics GmbH, Ebersberg, Germany |
| Pr7 | S1_*ZWF1_*fwd | AGTAAATCCAATAGAATAGAAAACCACATAAGGCAAGATGcgtacgctgcaggtcgac | Deletion of *ZWF1* in BY4742 | Eurofins Genomics GmbH, Ebersberg, Germany |
| Pr8 | S2_*ZWF1_*rev | AGTGACTTAGCCGATAAATGAATGTGCTTGCATTTTTCTAatcgatgaattcgagctcg | Deletion of *ZWF1* in BY4742 | Eurofins Genomics GmbH, Ebersberg, Germany |
| Pr9 | S1_*TRX1_*fwd | TTAGTGTAATAGAAGACTAGACACCTCGATACAAATAATGcgtacgctgcaggtcgac | Deletion of *TRX1* in BY4742 | Eurofins Genomics GmbH, Ebersberg, Germany |
| Pr10 | S2_*TRX1_*rev | CAGTATAGAAACACAATATATCGGTCATTGGGTGAGTTTAatcgatgaattcgagctcg | Deletion of *TRX1* in BY4742 | Eurofins Genomics GmbH, Ebersberg, Germany |
| Pr11 | S1_*TRX2_*fwd | ACGAGAGTCTACGATATCTTTAAATAACACATCAATAATGcgtacgctgcaggtcgac | Deletion of *TRX2* in BY4742 | Eurofins Genomics GmbH, Ebersberg, Germany |
| Pr12 | S2_*TRX2_*rev | ACATGATGTACTTTACGTAGCGTTAATATACCGGCAACTAatcgatgaattcgagctcg | Deletion of *TRX2* in BY4742 | Eurofins Genomics GmbH, Ebersberg, Germany |
| Pr13 | S1_*HIS3_*fwd | ACTAAAAAATGAGCAGGCAAGATAAACGAAGGCAAAGATGcgtacgctgcaggtcgac | Deletion of *HIS3* in CEN.PK113-1A | Merck KGaA, Darmstadt, Germany |
| Pr14 | S2_*HIS3_*rev | ATATATCGTATGCTGCAGCTTTAAATAATCGGTGTCACTAatcgatgaattcgagctcg | Deletion of *HIS3* in CEN.PK113-1A | Merck KGaA, Darmstadt, Germany |

**Supplementary Table 7: Yeast strains used in this study.**

| **Genotype** | **Source** |
| --- | --- |
| BY4742 *MAT*α *his3*∆*1 leu2*∆*1 lys2*∆*0 ura3*∆*0* | Euroscarf |
| BY4742 +p413TEF empty | This study |
| BY4742 +p413TEF PeredoxDS | This study |
| BY4742 +p413TEF NAPstar1 | This study |
| BY4742 +p413TEF NAPstar2 | This study |
| BY4742 +p413TEF NAPstar3 | This study |
| BY4742 +p413TEF NAPstar6 | This study |
| BY4742 +p413TEF NAPstar7 | This study |
| BY4742 +p413TEF NAPstarC | This study |
| BY4742 +p413TEF roGFP2-Grx1 | This study |
| BY4742 +p413TEF NERNST | This study |
| BY4742 ∆*zwf1*::*natNT2* | This study |
| BY4742 ∆*zwf1* +p413TEF empty | This study |
| BY4742 ∆*zwf1* +p413TEF PeredoxDS | This study |
| BY4742 ∆*zwf1*  +p413TEF NAPstar1 | This study |
| BY4742 ∆*zwf1* +p413TEF NAPstar2 | This study |
| BY4742 ∆*zwf1*  +p413TEF NAPstar3 | This study |
| BY4742 ∆*zwf1*  +p413TEF NAPstar6 | This study |
| BY4742 ∆*zwf1*  +p413TEF NAPstar7 | This study |
| BY4742 ∆*zwf1*  +p413TEF NAPstarC | This study |
| BY4742 ∆*glr1*::*kanMX4* | Morgan et al., 2013 |
| BY4742 ∆*glr1*  +p413TEF empty | This study |
| BY4742 ∆*glr1*  +p413TEF PeredoxDS | This study |
| BY4742 ∆*glr1*  +p413TEF NAPstar3 | This study |
| BY4742 ∆*glr1*  +p413TEF NAPstarC | This study |
| BY4742 ∆*glr1*  +p413TEF roGFP2-Grx1 | This study |
| BY4742 ∆*glr1*  +p413TEF NERNST | This study |
| BY4742 *trx1*::*kanMX4* ∆*trx2*::*natNT2* | This study |
| BY4742 +p413TEF empty | This study |
| BY4742 +p413TEF PeredoxDS | This study |
| BY4742 +p413TEF NAPstar3 | This study |
| BY4742 +p413TEF NAPstarC | This study |
| CEN.PK113-1A | Kind gift from P. Kötter, Frankfurt |
| CEN.PK113-1A∆*his3::hphNTI* | This study |
| CEN.PK113-1A∆*his3*  +p413TEF NAPstar4.3 | This study |
| CEN.PK113-1A∆*his3*  +p413TEF Peredox | This study |
| CEN.PK113-1A∆*his3*  +p413TEF HyPer7 | This study |
